# QTG-Finder2: a generalized machine-learning algorithm for prioritizing QTL causal genes in plants

**DOI:** 10.1101/2020.02.03.931444

**Authors:** Fan Lin, Elena Z. Lazarus, Seung Y. Rhee

**Author notes:** Corresponding author: Seung Y. Rhee, Department of Plant Biology, Carnegie Institution for Science, Stanford, California 94305, USA.

## Abstract

Linkage mapping has been widely used to identify quantitative trait loci (QTL) in many plants and usually requires a time-consuming and labor-intensive fine mapping process to find the causal gene underlying the QTL. Previously, we described QTG-Finder, a machine-learning algorithm to rationally prioritize candidate causal genes in QTLs. While it showed good performance, QTG-Finder could only be used in Arabidopsis and rice because of the limited number of known causal genes in other species. Here we tested the feasibility of enabling QTG-Finder to work on species that have few or no known causal genes by using orthologs of known causal genes as training set. The model trained with orthologs could recall about 64% of Arabidopsis and 83% of rice causal genes when the top 20% ranked genes were considered, which is similar to the performance of models trained with known causal genes. We further extended the algorithm to include polymorphisms in conserved non-coding sequences and gene presence/absence variation as additional features. Using this algorithm, QTG-Finder2, we trained and cross-validated *Sorghum bicolor* and *Setaria viridis* models. The *S. bicolor* model was validated by causal genes curated from the literature and could recall 70% of causal genes when the top 20% ranked genes were considered. In addition, we applied the *S. viridis* model and public transcriptome data to prioritize a plant height QTL and identified 13 candidate genes. QTL-Finder2 can accelerate the discovery of causal genes in any plant species and facilitate agricultural trait improvement.

## Introduction

Improving crop production to address the rapid increase in the global food demand, combined with increasing limitations of arable land, remains a major challenge. The world population is expected to exceed 9 billion by 2050 and will require a 70% increase in global food production (FAO 2009). Between 1985 and 2005 the world’s croplands increased by only about 2.4% (Foley *et al.* 2011). Therefore a significant increase of crop yield is required to feed the growing population, especially when there are increasing uncertainties of the changing climate.

Two common approaches used to improve crop yield and other agriculturally important traits are plant breeding and genome editing. Although plant breeding contributed significantly to crop yield improvement in the past century, it is facing obstacles such as limited sources of genetic variation and time-consuming phenotypying and germplasm evaluation (Rodriguez-Leal *et al.* 2017). With the advancement of CRISPR/Cas9 technology, genome editing has become a much faster way to enhance crop traits and it is possible to introduce novel alleles for a single gene via targeted mutagenesis (Rodriguez-Leal *et al.* 2017). However, genome editing will require identification of the trait-associated genes or the causal variants (Ramstein *et al.* 2019). Many trait-associated causal genes in quantitative trait loci (QTL) have been validated by mutational analysis and functional complementation experiments (Weigel and Nordborg 2005).

As one of the most commonly used genetic mapping methods, thousands of QTL mapping studies have been conducted on many crops (Yonemaru *et al.* 2010; Mace *et al.* 2019) but the causal genes for most of these QTLs have not been identified or validated by experiments. For example, there are less than one hundred curated causal genes that have been cloned and validated by complementation experiments in Arabidopsis or rice, and there are even fewer known causal genes in other plant species (Martin and Orgogozo 2013). Identifying causal genes from hundreds to thousands of genes in a QTL region usually requires a great amount of time and effort to fine map the causal gene (Huang *et al.* 2016a). Therefore, a computational method to predict or prioritize causal genes will be helpful for accelerating the discovery of novel trait-associated genes in QTLs.

Previously we developed a machine-learning based algorithm, named QTG-Finder, to prioritize causal genes in QTLs (Lin *et al.* 2019). The algorithm uses additional information such as polymorphisms from re-sequencing data, function annotation, co-function network, gene essentiality and paralog copy number to prioritize causal genes in QTLs. We trained models for *Arabidopsis thaliana* (Arabidopsis) and *Oryza sativa* (rice) with curated causal genes in each species. Based on validation using an independent set of newly curated genes, the models could recall about 64% of Arabidopsis and 79% of rice causal genes when the top 20% ranked genes in a QTL were considered. However, the models were only developed for Arabidopsis and rice and were trained on a relatively small number of known causal genes from each species. There were insufficient numbers of known causal genes in other plants to develop such predictive models.

To devise an algorithm that would work even on species with few or no training data available, we wondered whether orthology could be used to create or extend training data. This idea was based on several factors. First, many causal genes identified by linkage mapping are evolutionary hotspots (Martin and Orgogozo 2013). Second, in plants and animals, some genes have repeatedly been major components of phenotypic variation of similar traits (Gompel and Prud’homme 2009; Kopp 2009). For example, *Flowering Time* (*FT*) has been reported to be a causal gene of flowering time QTLs in Arabidopsis (Kojima *et al.* 2002; Schwartz *et al.* 2009), barley (Yan *et al.* 2006), wheat (Yan *et al.* 2006), sunflower (Blackman *et al.* 2010) and ryegrass (Skot *et al.* 2011). There are many other examples of conservation in causal genes for the same trait across species (Martin and Orgogozo 2013). Therefore, we hypothesized that the orthologs of causal genes are also likely to be causal genes.

We tested this hypothesis by training models in Arabidopsis and rice with orthologs of known causal genes. The performance indicated the feasibility of this approach. We further tested the approach by training models for *Sorghum bicolor* (sorghum) and *Setaria viridis* (Setaria), which have only few known causal genes. We validated the sorghum model by testing QTLs with known causal genes curated from the literature. We also demonstrated the usage of the Setaria model by combining the prioritization results with published transcriptome data to obtain 13 causal gene candidates for a Setaria height QTL.

## Materials and Methods

### The orthologs of known causal genes

The list of causal genes used for orthology analysis was based on a list of causal alleles previously published (Martin and Orgogozo 2013). Since the original list only provided the gene name of those causal genes, we first curated their gene ID or UniProt ID from the references cited. When the ID was not available in the papers, we searched the gene name in genome annotation databases such as RAP-DB (https://rapdb.dna.affrc.go.jp), maizeGDB (https://www.maizegdb.org), soyKB (http://soykb.org/) or the UniProt database (https://www.uniprot.org). The gene ID or UniProt ID was used as a query to search the EggNOG database (v4.5.1) (Huerta-Cepas *et al.* 2016) to obtain the ortholog group to which it belongs and its fine-grained orthologs. Fine-grained orthologs in EggNOG is defined as orthologs derived from a pairwise orthology between members of two species in an orthologous group based on phylogenic analysis. For genes that were not found in EggNOG, we obtained their protein sequences from UniProt or Genbank and used a HMMER-based sequence search (http://eggnogdb.embl.de/#/app/seqscan) to find the ortholog group. When available, the fine-grained orthologs were used as the ortholgs of causal genes. When fine-grained orthologs were not available, all members in the ortholog group were used as orthologs. We examined the orthology in major crops and model organisms of eudicots and monocots: *Arabidopsis thaliana*, *Solanum lycopersicum*, *Brassica rapa*, *Glycine max*, *Oryza sativa japonica*, *Oryza sativa indica*, *Setaria italica*, *Sorghum bicolor*, *Brachypodium distachyon*, *Hordeum vulgare* and *Zea mays* (Supplemental Table S1).

We obtained the ortholog list for *Setaria viridis* in a different way because the EggNOG database only includes *S. italica* genes and not *S. viridis* genes. Since *S. italica* is a domesticated line derived *from S. viridis* and has excellent collinearity with *S. viridis* (Supplemental Figure S1, (Bennetzen *et al.* 2012), we used DAGChainer (Haas *et al.* 2004) in CoGe to identify collinear gene pairs that fall in contiguous chains between *S. viridis* and *S. italica*. The DAGChainer results allowed us to convert *S. italica* gene IDs to *S. viridis* gene IDs (Supplemental Table S1).

### Building new features based on polymorphisms in conserved non-coding regions and gene presence/absence for Arabidopsis and rice models

The conserved non-coding regions, Conserved Elements (CE) and Transcription Factor Binding Sites (TFBS), were downloaded from the Plant Transcriptional Regulatory Map (PlantRegMap, last modified on 2019-10-11, http://plantregmap.cbi.pku.edu.cn/). The CEs were identified based on the genome alignments of plants (Jin *et al.* 2014). The TFBSs were based on the correlation between frequencies in binding motifs and conservation scores (Tian *et al.* 2019). The TFBSs and CEs were assigned to genes that were located within 1kb upstream or downstream of the genes. We incorporated the TFBSs and CEs as regulatory annotations in SnpEff (v 4.3r) and identified SNPs and Indels in these TFBSs or CEs. We counted the number of SNPs and Indels in these conserved non-coding regions for each gene and built four features: CE_snp, TFBS_snp, CE_indel, TFBS_indel.

The gene presence/absence data were obtained from published studies. The Arabidopsis gene presence/absence data were based on 80 *A. thaliana* accessions with 10 to 24x sequencing coverage (Tan *et al.* 2012). The rice gene presence/absence data were based on 453 *O. sativa* accessions with sequencing depth of over 20x (Hu *et al.* 2018). The percentage of absence across the sequenced accessions was calculated for each gene and used as the feature “percent_absence”.

### Features and model training of sorghum and Setaria models

Features for sorghum and Setaria models were generated as follows. Polymorphism features were extracted in the same way as previously described (Lin *et al.* 2019). Briefly, SNP data were annotated by SIFT4G (v 2.4) and SnpEff (v 4.3r) and assigned to each gene in the genome. The polymorphism features were mostly binary features that represent the presence of a specific type of SNP for each gene. For example, if a gene contained any deleterious non-synonymous SNPs, the “is_nonsyn_deleterious” feature was set to 1, otherwise it was set to 0. The sorghum SNP data were downloaded from Sorghum Genome SNP Database (SorGSD) (Luo *et al.* 2016), which provides SNP data for a diverse panel of 48 sorghum lines. The Setaria SNP data and gene presence/absence data are based on a diverse panel of 598 *S. viridis* accessions (Huang *et al.* 2019). Since there is no pre-built SnpEff database for *S. viridis*, we built a *S. viridis* database using the .gff file of *S. viridis* v2.1 downloaded from Phytozyme12 (https://phytozome.jgi.doe.gov).

The Gene Ontology (GO) annotations for sorghum and Setaria were obtained from PLAZA4.0 (https://bioinformatics.psb.ugent.be/plaza/versions/plaza_v4_monocots/) (Van Bel *et al.* 2018). We used GOslim (http://current.geneontology.org/ontology/subsets/goslim_metagenomics.obo) to aggregate the molecular function GO terms to higher-level terms such as is_transporter and is_transcription_factor. For genes encoding an enzyme, we further determined their metabolic domains based on Plant Metabolic Network databases (PMN, release 12.5) (Schlapfer *et al.* 2017).

Paralog copy number of each gene was determined by OrthoFinder (v2.3.3) (Emms and Kelly 2019). We used the DIAMOND algorithm and the default setting of OrthoFinder. Protein sequences of thirteen species or subspecies were downloaded from PLAZA4.0, including *Arabidopsis thaliana*, *Brachypodium distachyon*, *Brassica rapa*, *Glycine max*, *Hordeum vulgare*, *Oryza sativa japonica*, *Oryza sativa indica*, *Populus trichocarpa*, *Sorghum bicolor*, *Setaria viridis*, *Setaria italica, Solanum lycopersicum* and *Zea mays*. Paralogs were counted for each species in the orthologous group determined by OrthoFinder.

The Setaria and sorghum models were trained with orthologs of causal genes from any other plant species. We used the random forest algorithm, which is an ensemble learning method that fits a number of decision trees on various sub-sampled datasets (Tin Kam 1998). Random forest integrates the votes from these trees to improve accuracy and reduce the chance of over-fitting. Model parameters including the number of trees, maximum number of features to consider when seeking for the best split, the minimum number of samples required to split a node in a decision tree and the ratio of positives and negatives in the training set were used to optimize the models to maximize cross-validation AUC-ROC (Supplemental Figures S2 and S3).

### Cross-validation, external validation and feature importance analysis

The methods for cross-validation, external validation and feature importance analysis were the same as previously described (Lin *et al.* 2019). The causal genes used for external validation were not used for training the QTG-Finder models. However, some of them are orthologs of known causal genes in other species, which could have been used for training QTG-Finder2 models. We therefore excluded these orthologs from the training set of QTG-Finder2 to avoid over-estimation of model performance.

The external validation of the sorghum model was conducted on a set of causal genes curated from the literature (Table 1) (Magalhaes *et al.* 2007; Jordan *et al.* 2010; Kawahigashi *et al.* 2011; Murphy *et al.* 2011; Lin *et al.* 2012; Saballos *et al.* 2012; Murphy *et al.* 2014; Yang *et al.* 2014; Boyles *et al.* 2017; Hilley *et al.* 2017). We applied the model to all genes in the QTL regions, which were defined by the literature.

**Table 1.** Curated *S. bicolor* QTL causal genes and external validation results.

| QTL Trait | Gene name | Gene ID | Genes in QTL | Percent rank | Reference |
| --- | --- | --- | --- | --- | --- |
| Light sensitivity | *phyB* | Sobic.001G394400 | 144 | 1% | Yang *et al.*, 2014 |
| Brown midrib | *bmr2* | Sobic.004G062500 | 27 | 3% | Saballos *et al.*, 2012 |
| Amylose | *Wx* | Sobic.010G022600 | 706 | 3% | Boyles *et al.*, 2017 |
| Fungus resistance | *Ds1* | Sobic.005G065000 | 389 | 15% | Kawahigashi *et al.*, 2011 |
| Plant height | *Dw2* | Sobic.006G067700 | 335 | 15% | Hilley *et al.*, 2017 |
| Seed shattering | *Sh1* | Sobic.001G199200 | 117 | 18% | Lin *et al.*, 2012 |
| Aluminum tolerance | *MATE* | Sobic.003G403000 | 26 | 19% | Magalhaes *et al.*, 2007 |
| Light sensitivity | *ghd7* | Sobic.006G004400 | 115 | 61% | Murphy *et al.*, 2014 |
| Pollen fertility | *PPR* | Sobic.002G057050 | 26 | 69% | Jordan *et al.*, 2010 |
| Flowering time | *PRR37* | Sobic.006G057900 | 22 | 77% | Murphy *et al.*, 2011 |

We used Fisher’s exact test for the pairwise comparison of external validation results. Each gene in the external validation set was assigned to one of two classes: (1) the gene was included in the prioritized fraction (e.g. top 5%, 10% or 20%), or (2) the gene was not included in the prioritized fraction. The number of genes in the two classes (prioritized vs. not prioritized) was used for Fisher’s exact tests.

### Sequence alignment for candidate genes

Multiple sequence alignments were performed using Clustal Omega (v1.2.4, https://www.ebi.ac.uk/Tools/msa/clustalo/) across grass species including *Brachypodium distachyon*, *Panicum virgatum, Oryza sativa*, *Setaria viridis*, *Setaria italica, Sorghum bicolor* and *Zea mays*. Homologous protein sequences in these species were obtained from Phytozyme12. Human and yeast RIO2 sequences were obtained from NCBI (https://www.ncbi.nlm.nih.gov).

For promoter sequence comparison, we used pairwise global sequence alignment (EMBOSS Needle, https://www.ebi.ac.uk/Tools/psa/emboss_needle/). The promoter sequences were defined as 1kb upstream of coding sequence (CDS) and downloaded from Phytozome12. We further examined the putative Transcription Factor Binding Sites (TFBS) in the promoters. TFBSs were predicted by Plant Transcriptional Regulatory Map tool (PlantRegMap, http://plantregmap.cbi.pku.edu.cn/binding_site_prediction.php).

### Data Availability

The source code and training data for Arabidopsis, rice, sorghum and Setaria are available at Github (https://github.com/carnegie/QTG_Finder). The pre-trained QTG2-Finder2 models are available at Amazon cloud storage through the links provided below. All models were trained and tested using Python 3.7.3 and scikit-learn 0.21.2. https://carnegiedpb.s3.amazonaws.com/software/QTG2_prediction/AT_model.dat.zip https://carnegiedpb.s3.amazonaws.com/software/QTG2_prediction/OS_model.dat.zip https://carnegiedpb.s3.amazonaws.com/software/QTG2_prediction/SB_model.dat.zip https://carnegiedpb.s3.amazonaws.com/software/QTG2_prediction/SV_model.dat.zip

All supplemental materials are available at FigShare.

## Results

### Incorporating an orthology approach in the QTG-Finder2 algorithm

The QTG-Finder algorithm described previously only used known causal genes of a single species to train a model of that species (Figure 1A). For QTG-Finder2, we trained models with not only the known causal genes in the target species but also the orthologs of causal genes from other species (Figure 1B).

**Figure 1.**
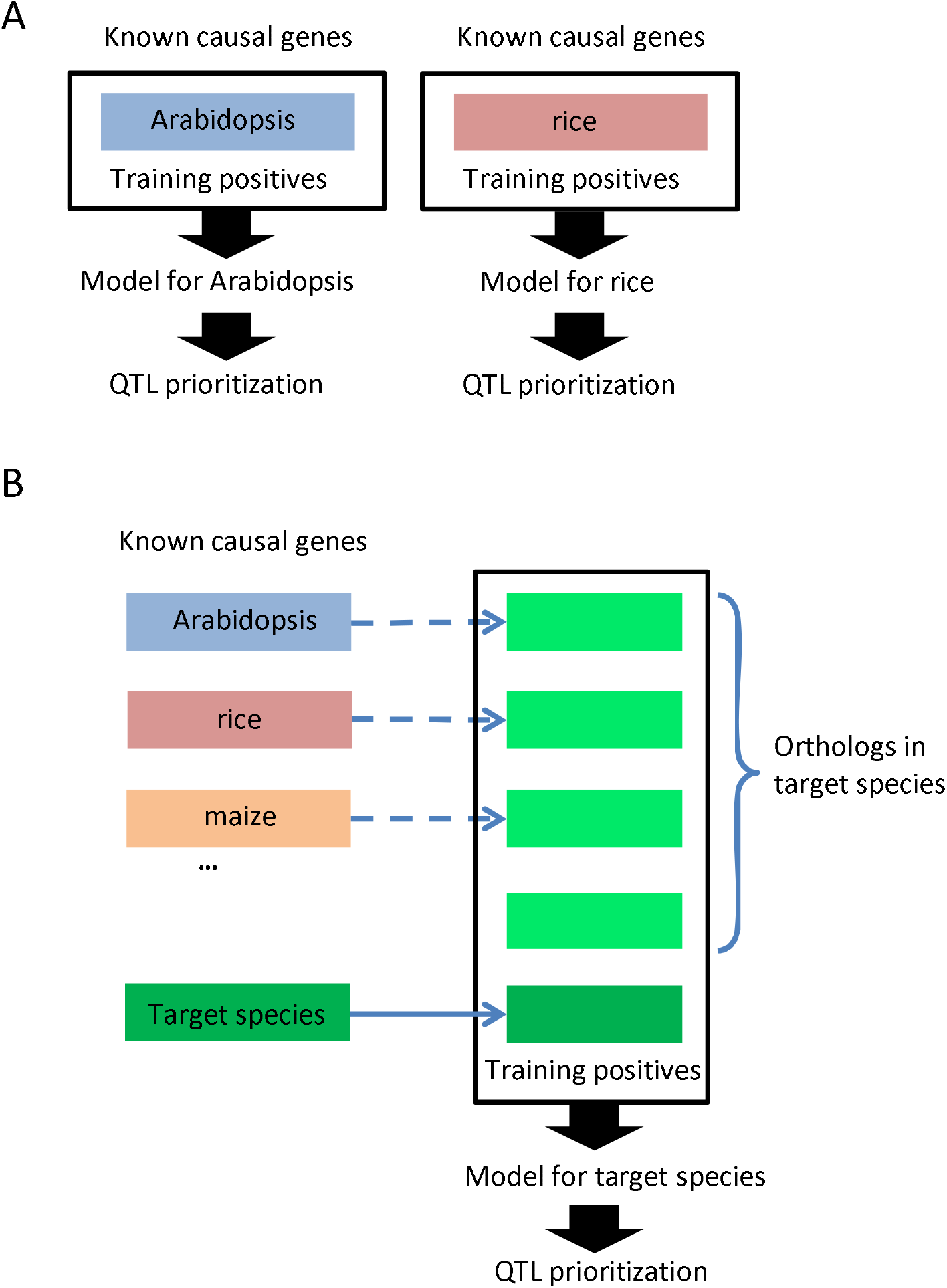
Incorporating an orthology approach to the QTG-Finder algorithm facilitates training models in other plant species (A) The original QTG-Finder algorithm. Only the known causal genes were used to train a model for a given species. (B) QTG-Finder2 algorithm. Orthologs of the known causal genes from any species were also used to train a model. This method allows QTG-Finder to be implemented in species without enough or any known causal genes.

With 253 curated causal genes from any plant species (Figure 2), we identified their orthologs in 12 species and subspecies: *Arabidopsis thaliana*, *Solanum lycopersicum*, *Brassica rapa*, *Glycine max*, *Oryza sativa japonica*, *Oryza sativa indica*, *Setaria italica*, *Setaria viridis*, *Sorghum bicolor*, *Brachypodium distachyon*, *Hordeum vulgare* and *Zea mays* (Supplemental Table S1). The 12 species included major crops and model organisms of eudicots and monocots. Each species had orthologs for about 60% of the causal genes (Supplemental Figure S4A). Some causal genes had multiple orthologs and the average number of orthologs varied across species (Supplemental Figure S4B).

**Figure 2.**
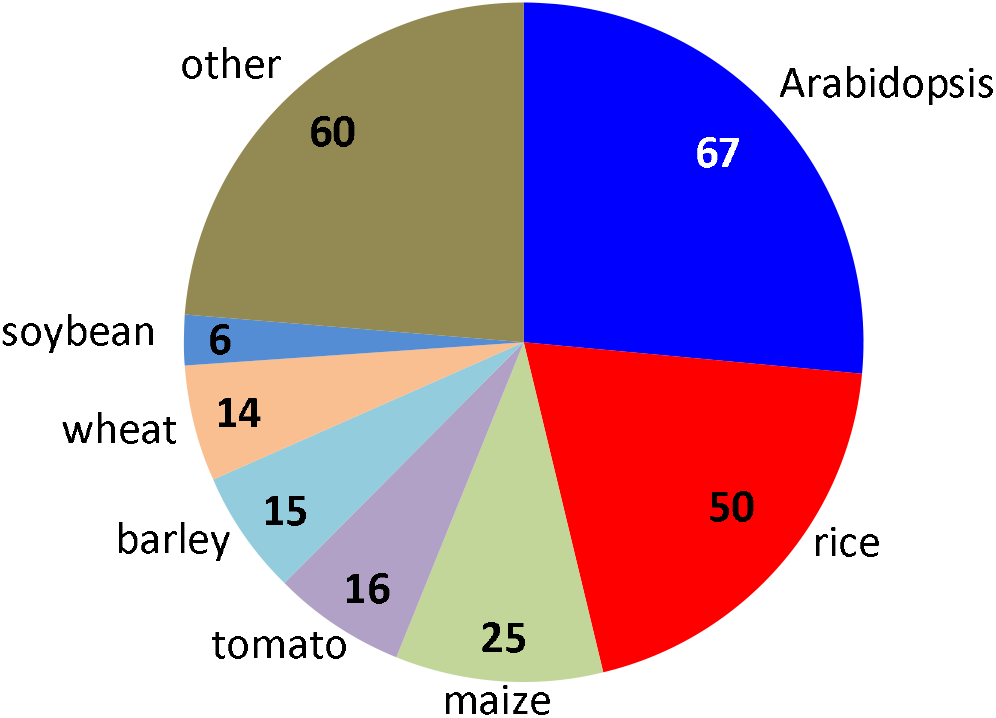
Most plant species do not have enough curated known causal genes to train a model as Arabidopsis or rice does. Numbers indicate the number of known causal genes in each species. Arabidopsis, *Arabidopsis thaliana*; rice, *Oryza sativa japonica;* maize, *Zea mays*; tomato, *Solanum lycopersicum*; barley, *Hordeum vulgare*; wheat, *Triticum aestivum*; soybean, *Glycine max*. Data from Martin and Orgogozo, 2013

### Testing Arabidopsis and rice models trained with orthologs

We asked if the models trained with orthologs would perform as well as models trained with only the known causal genes in the target species. To test this hypothesis, we trained models in Arabidopsis and rice using three different sets of positive training data: 1) only known causal genes in the target species, 2) only orthologs, and 3) known causal genes plus orthologs. For the Arabidopsis model, we used 60 known causal genes from Arabidopsis and 146 orthologs of causal genes from other species (Figure 3A). In the rice model, we used 45 known causal genes from rice and 206 orthologs of causal genes from other species.

**Figure 3.**
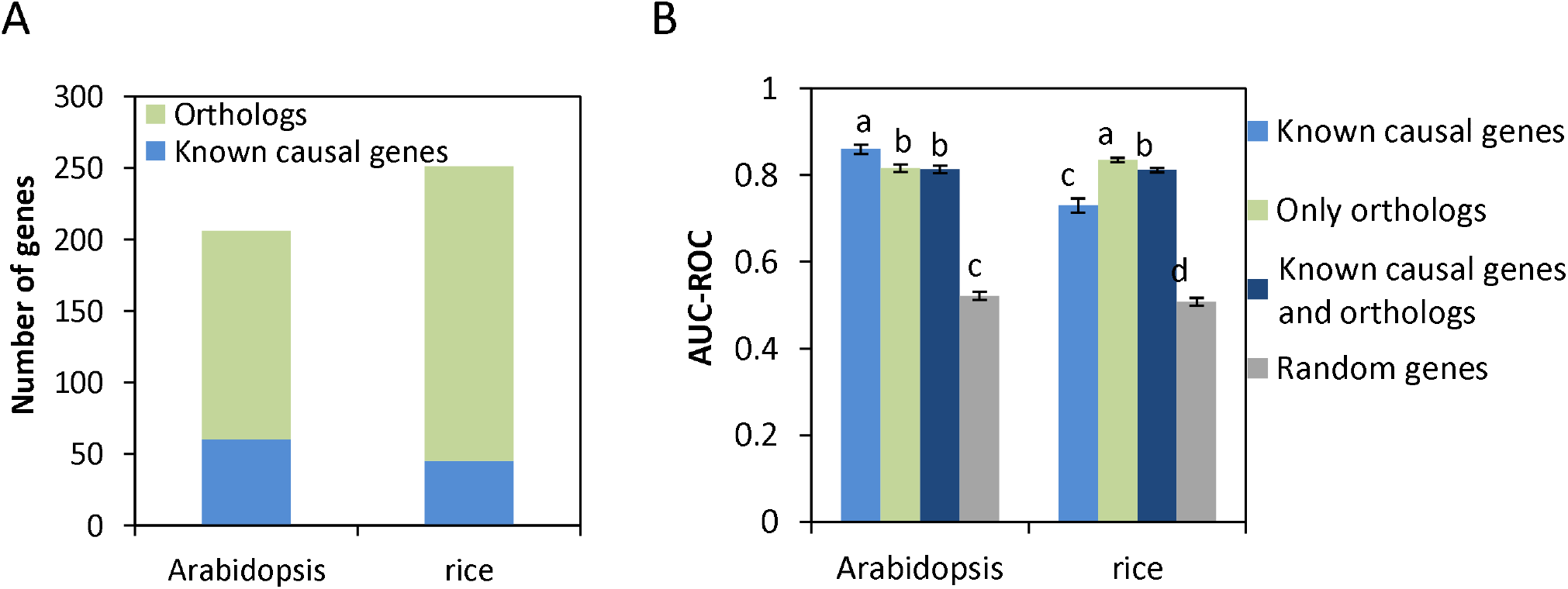
Models trained with orthologs have comparable performance as the models trained with known causal genes (A) The number of known causal genes and orthologs derived from causal genes in any plant species (B) Cross-validation of models trained with known causal genes, with orthologs, or with causal genes plus orthologs. AUC-ROC (Area Under the Curve - Receiver Operating Characteristic) was used to compare training performance of the models. Error bars represent standard deviation, N=50 for each bar. One-way ANOVA followed by Tukey HSD post-hoc test was performed to determine the statistical difference (p<0.05) among the groups as represented by letters.

We first performed cross-validation to evaluate models that were trained with the three training sets (Figure 3B). We used the Area Under the Receiver Operating Characteristic Curve (AUC-ROC) to evaluate the training performance of these models. For both species, models trained with any of the training sets was significantly higher than the models trained with randomly selected genes (Figure 3B, One-way ANOVA followed by Tukey HSD post-hoc test, p-value < 0.05). In Arabidopsis, the models trained with only the known causal genes had the highest average AUC-ROC score (0.86). The model trained with only orthologs had an average AUC-ROC of 0.82, which is comparable to the model trained with known causal genes. The model trained on both the orthologs and known causal genes had an average AUC-ROC of 0.81, which was not distinguishable from the model with just the orthologs. This indicates that orthologs have similar properties as known causal genes. Compared to these scores, the model trained with randomly selected genes had an average AUC-ROC of 0.52. In rice, the model trained with only orthologs had the highest average AUC-ROC of 0.84. This is not simply due to the sample size increasing since this trend was not observed in Arabidopsis where the sample size was also increased in the ortholog training data. Interestingly, the model trained with only the known genes showed the lowest score of 0.73. Combining the orthologs with the known causal genes for training gave a score of 0.81. Since the models trained with only orthologs had significantly higher AUC-ROC than models trained with random genes, these results indicate the orthologs by themselves will be useful for model training.

After optimizing model parameters from cross-validation results, we evaluated the final models with an external validation set, which contained independently curated causal genes that had not been seen by the models. The validation method and data set were the same as previously described (Lin *et al.* 2019). For each causal gene in the validation set, the models were applied to rank all genes in the QTL region where the causal gene is located. We applied the model to rank and prioritize the top 5%, 10% and 20% of genes in the QTL region and examined if the known causal gene was included in the prioritized gene list. All models performed significantly better than the models trained with randomly selected genes (Figure 4). The models trained with only orthologs not only performed significantly better than background, but also were not different from models trained with just the known causal genes. For Arabidopsis, the model trained with only orthologs could recall 27%, 36% and 64% of causal genes at the top 5%, 10% and 20% cutoffs, respectively (Figure 4). For rice, the model trained with only orthologs can recall 28%, 56% and 83% of causal genes at the top 5%, 10% and 20% cutoffs, respectively (Figure 4). At all three cut-offs, the Arabidopsis models that combined the Arabidopsis causal genes and orthologs performed better than models using either causal genes or orthologs alone, though not significantly (Fisher’s exact test, p-value > 0.05). For rice, all three models performed similarly. All the Arabidopsis and rice models performed significantly better than the background at 10% and 20% cutoffs (Fisher’s exact test, p-value<0.05) but not at 5% cutoff. The background was determined as the theoretical probability of including the causal gene when we randomly selected 5%, 10% or 20% of the genes from the QTL region.

**Figure 4.**
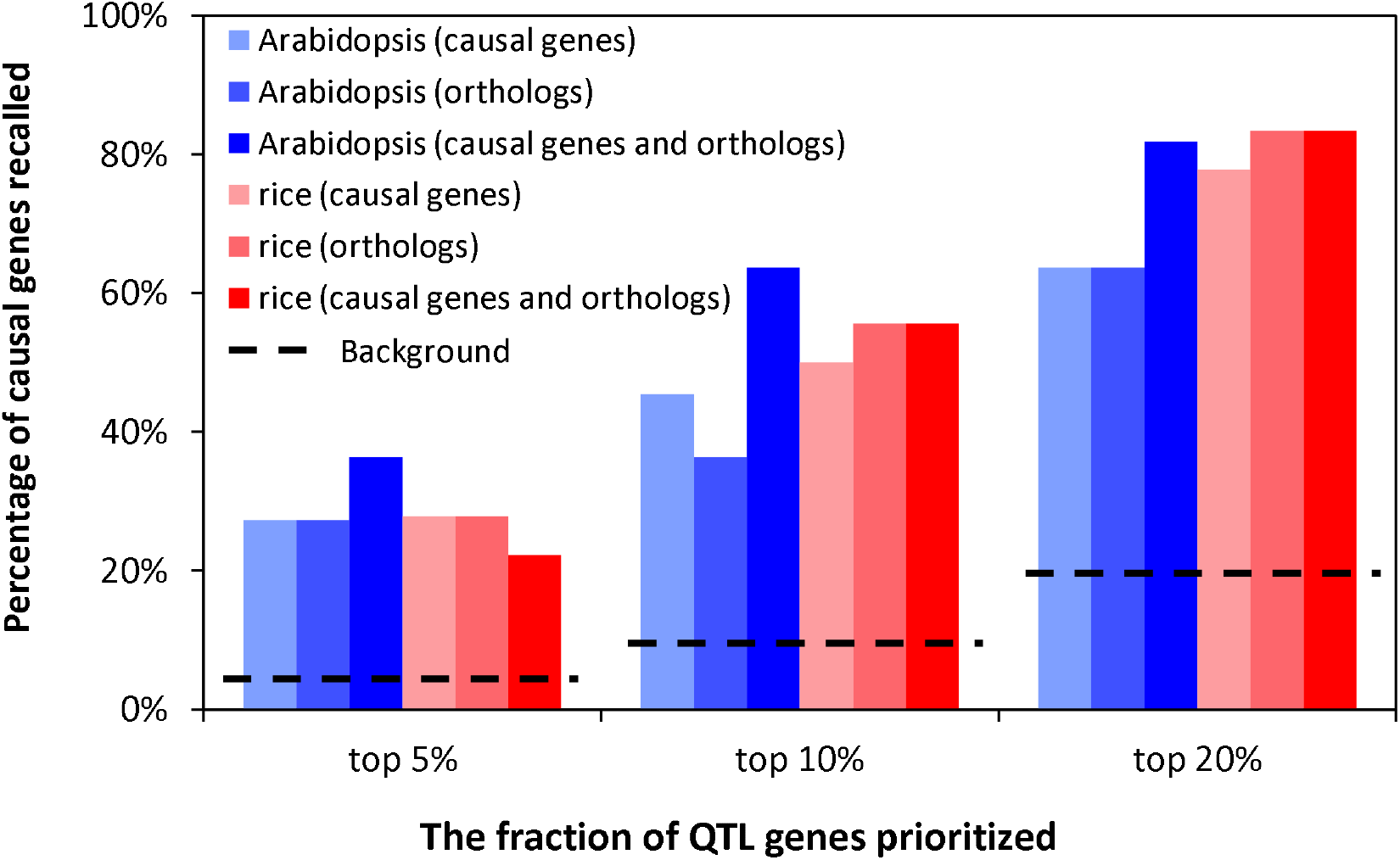
Models trained with known causal genes and orthologs have similar performance based on external validation. Model performance was evaluated when the top 5%, 10% or 20% of the ranked QTL genes were considered. Fisher’s exact test was performed between the models trained only with known causal genes versus the models trained with only orthologs or the models trained with causal genes plus orthologs. No statistical difference was detected (p>0.05). The black dashed lines indicate theoretical background estimated by the percentage of causal genes being recalled when we randomly selected 5%, 10% or 20%of the genes from the QTL region.

To determine if different levels of orthology affected performance, we compared the orthology method described above with two alternative methods: 1) using taxon-constrained orthologs and 2) using only EggNOG’s fine-grained orthologs defined as orthologs derived from a pairwise orthology between members of two species in an orthologous group based on phylogenic analysis (Huerta-Cepas *et al.* 2016). For the taxon-derived orthology method, we only considered the othologs for species that are in the same lineage (monocot or eudicot) as the causal gene being queried. For example, if the known causal gene were identified in a monocot species, then we would only consider its orthologs in monocot species. Using the same external validation set, we compared the original orthology method with these two methods and found that their performance was similar to each other (Fisher’s exact test, p-value>0.05) (Supplemental Figure S5).

Models trained with both the known causal genes in the species and orthologs from other species represent a more generalized model since it combines information from known causal genes in target species and information from causal genes in other species. The models trained with known causal genes plus orthologs performed similarly as the models trained with only orthologs in cross-validation and external validation (Figures 3 and 4). Therefore, we used models trained with known causal genes plus orthologs for subsequent analyses.

### Exploring and adding new features to QTG-Finder2

We explored new features that may help distinguish causal genes from other genes such as polymorphisms in conserved non-coding regions and structural variations such as gene presence/absence. SNPs or Indels in some conserved non-coding sequences may disrupt transcription factor binding and influence gene expression patterns and traits. In addition, gene presence/absence variation has been linked to phenotypic variations. For example, causal genes like *RLM3* and *FRIGIDA* in Arabidopsis (Werner *et al.* 2005; Staal *et al.* 2008), *Sub1A* in rice (Xu *et al.* 2006) and *ZCCT1* and *ZCCT2* in barley (Yan *et al.* 2004) are absent in some accessions.

To generate features from polymorphisms in non-coding sequences, we used two types of predicted non-coding sequences in PlantRegMap (Tian *et al.* 2019): Conserved Elements (CE) and functional Transcription Factor Binding Sites (TFBS). In Arabidopsis and rice, there are significantly more SNPs and Indels in the CEs nearby causal genes than in the CEs nearby an average gene in the genome (Mann-Whitney U Test, p-value <0.05, Figure 5A, Supplemental Tables S2 and S3). However, the SNPs or Indels in TFBS were not significantly different between causal genes and non-causal genes (Mann-Whitney U Test, p-value >0.05, Figure 5A, Supplemental Tables S2 and S3).

**Figure 5.**
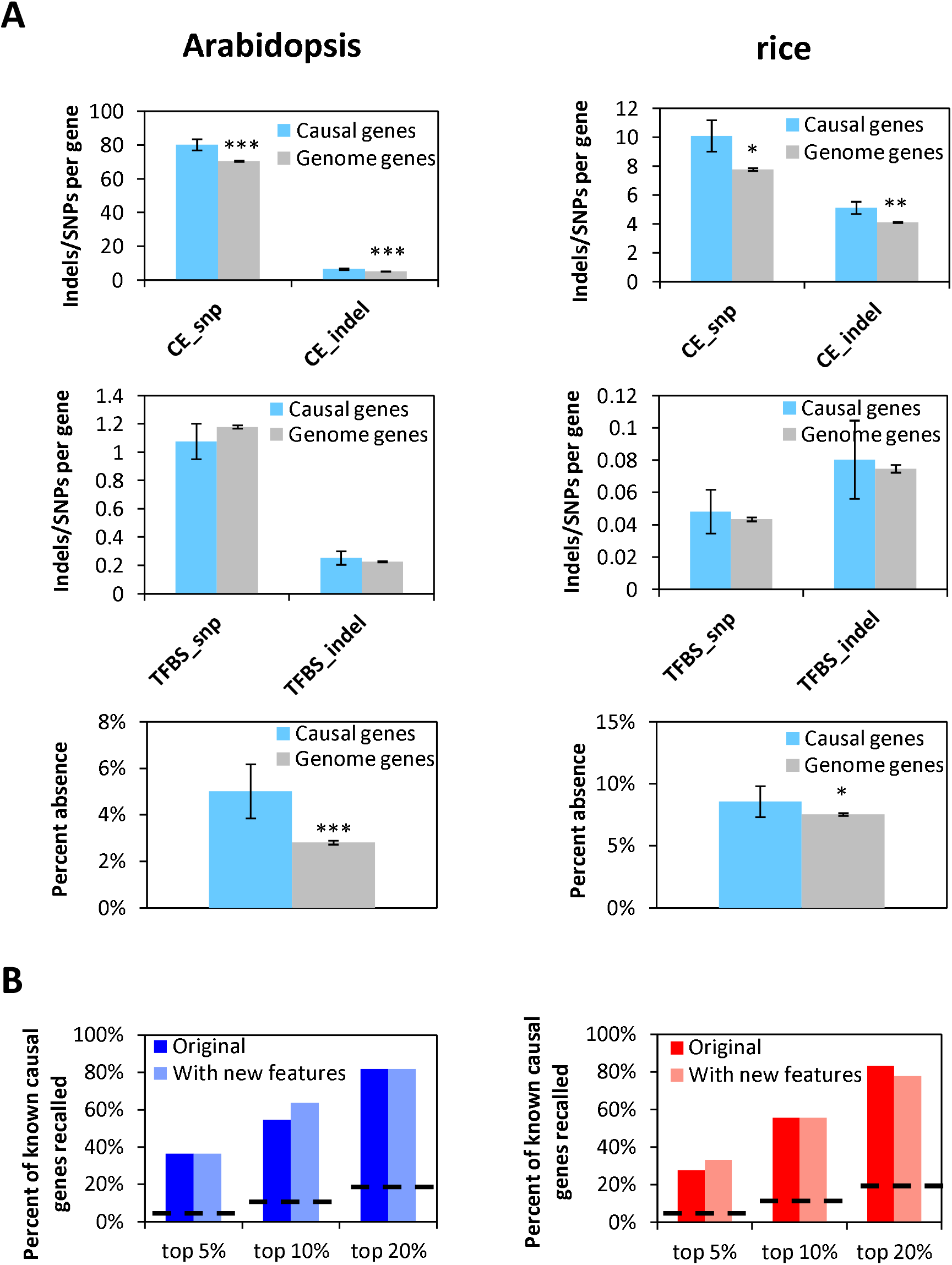
Exploring new features for QTG-Finder2 (A) Several new features we added were enriched for causal genes relative to an average genome gene. New features include the number of SNPs and Indels in conserved non-coding sequences flanking genes and the percent absence of a gene in the collection of natural variant accessions. Mann-Whitney U Test was used to compare the statistical difference between causal genes and genome genes. Significance levels were defined as: *, p<0.05, **, p<0.01,***, p<0.001. (B) Model performance does not change much after adding new features based on external validation. Independent validation sets were used to evaluate model performance. The black dashed lines indicate theoretical background. Fisher’s exact test was performed between models with and without new features. No statistical difference was detected (p>0.05).

In addition, we constructed a percent absence feature using previously published gene presence/absence analyses (Tan *et al.* 2012; Hu *et al.* 2018) In both Arabidopsis and rice, the causal genes had significantly higher percent absence than genome genes (Mann-Whitney U Test, p-value <0.05, Figure 5A, Supplemental Tables S2 and S3).

We were encouraged by the enrichment of the CEs and presence/absence variation in the causal genes and added them as new features. However, these new features did not change the model performance significantly (Fisher’s exact test, p-value>0.05). We first compared the external validation results for models with or without the new features (Figure 5B). Then, we examined the feature importance of those new features by using a leave-one-out analysis (Supplemental Figure S6). The CE_snp feature was the fourth most important feature in the Arabidopsis model. The other new features were not within the top 5 most important features. Since these new features do not reduce model performance, we kept them in the algorithm for subsequent analyses.

### Applying QTG-Finder2 to train sorghum and Setaria models

To test whether the QTG-Finder2 algorithm can be used to train models for species that have few or no known causal genes, we trained models in *Setaria viridis* (Setaria) and *Sorghum bicolor* (sorghum) with the orthologs derived from causal genes in other species. *S. bicolor* is an important C4 photosynthesis crop with excellent drought resistance (Calvino and Messing 2012). *S. viridis* is a C4 photosynthesis model grass and the wild ancestor of foxtail millet (*Setaria italica*), an important crop in Asia and Africa (Huang *et al.* 2016b).

We first conducted cross-validation for the Setaria and sorghum models (Figure 6A). The AUC-ROCs were 0.79 (Setaria model) and 0.77 (sorghum model), respectively, which were similar to the Arabidopsis and rice models trained on causal genes.

**Figure 6.**
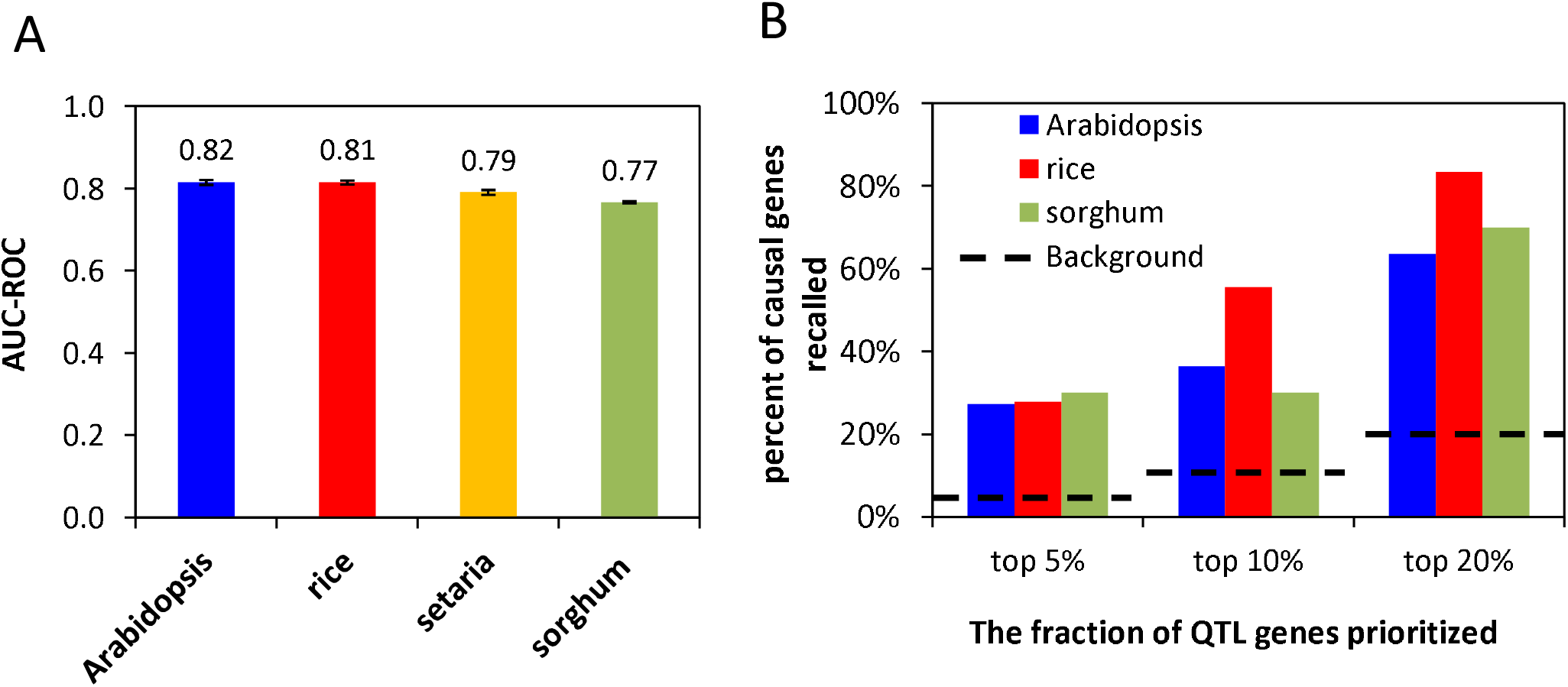
Performance of Setaria and sorghum models (A) The new *Sorghum bicolor* (sorghum) and *Setaria viridis* (Setaria) models trained by QTG-Finder2 algorithm have similar performance as the *Arabidopsis thaliana* (Arabidopsis) and *Oryza sativa* (rice) models. Cross-validation indicated by AUC-ROC (Area Under the Curve - Receiver Operating Characteristic). Error bars indicate standard deviation, N=50 per species. (B) External validation of the models for Arabidopsis, rice and sorghum that had independent causal gene data available. Fisher’s exact test was performed between the sorghum model versus Arabidopsis or rice model, and no statistical difference was detected (p>0.05). The black dashed lines indicate the theoretical background when the same fraction of genes in the QTL was randomly prioritized.

To validate the sorghum model, we performed an independent external validation. We curated ten sorghum causal genes from the literature (Table 1). When the top 5%, 10% and 20% of the genes in the QTL region were prioritized by the sorghum model, 30%, 30% and 70% of the causal genes were recalled, respectively (Figure 6B). The sorghum model’s performance was similar to the Arabidopsis model, which recalled 27%, 36% and 64% of the causal genes when the top 5%, 10% and 20% of the genes in the QTL were prioritized. While the performance of the sorghum model was lower than that for the rice model, there was no statistical difference in the performance between sorghum and Arabidopsis or rice models. All models performed significantly better than the background where the same number of genes were randomly prioritized at 10% and 20% cutoffs (Fisher’s exact test, p-values>0.05, Figure 5B) but not at 5% cutoff.

### Combining the Setaria model and transcriptome data to prioritize causal genes for a Setaria height QTL

Since there are no cloned QTL causal genes available for Setaria, we could not evaluate Setaria model’s performance with independent data. To demonstrate the usage of the Setaria model, we applied it to prioritize a well-determined plant height QTL in Setaria. This QTL is located on chromosome 5 and has been reported by two independent studies (Mauro-Herrera and Doust 2016; Feldman *et al.* 2017) and has large effects on height under many conditions such as different watering levels and density of planting (Feldman *et al.* 2017). There are 335 genes in the LOD1.5 interval of this major QTL (Feldman *et al.* 2017).

To select a testable number of candidates for this height QTL, we combined the Setaria model with published transcriptome data (Martin *et al.* 2016). We first applied the Setaria model to the QTL and prioritized 67 genes that ranked within the top 20%. Given that the experimental validation for this number of candidates would still constitute a large effort at this time, we incorporated transcriptome data to further narrow down the candidate gene list. Based on a transcriptome study on the developing internode of *S. viridis*, we selected genes that were up-regulated by more than 2-fold in the internode meristem or cell elongation zone relative to the maturation zone. We posited that genes that were up-regulated in these zones are more likely to be involved in internode elongation and therefore contribute to plant height. In the QTL interval, 60 genes were up-regulated either in the meristem or elongation zone relative to the maturation zone (Figure 7A). By comparing the top 20% of the prioritized genes with the up-regulated genes, we found 13 genes that met both criteria (Figure 7B, Supplemental Table S4).

**Figure 7.**
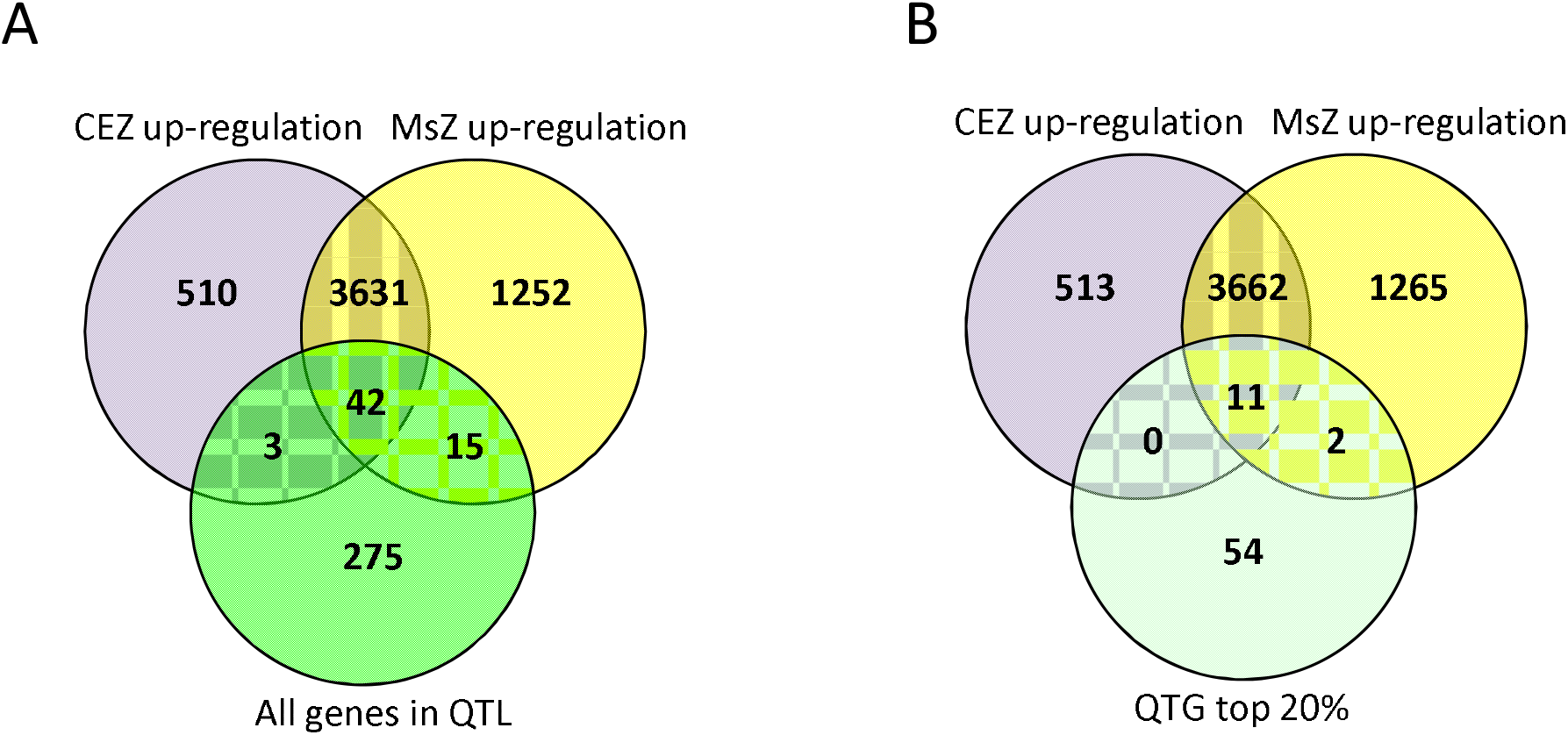
Candidate causal genes of a Setaria plant height QTL prioritized by QTG-Finder2 and transcriptome analysis (A) The overlap among genes up-regulated in the Meristem Zone (MsZ) and the Cell Elongation Zone (CEZ) relative to the maturation zone of the internode and all genes in the height QTL interval. (B) The overlap among genes up-regulated in the Meristem Zone (MsZ) and the Cell Elongation Zone (CEZ) relative to the maturation zone of the internode and the top 20% genes prioritized by QTG-Finder2 (QTG top 20%). Transcriptome data were obtained from Martin, 2016.

In addition to the 13 candidates we prioritized, there is one gene (*Semidwarf*, *SD1*, Sevir.5G410400) in this QTL interval, which was suggested to be a putative causal gene, though it has not been experimentally validated. *SD1* gene encodes gibberellin20 oxidase2 in rice, involved in gibberellin biosynthesis, and a loss of function allele gives dwarf phenotype in rice (Spielmeyer *et al.* 2002). The putative causal gene *SD1* has a percent rank of 24% according to the prediction of our Setaria model. Though not within the top 20% ranked genes, *SD1* is up-regulated in the meristem zone of *S. viridis* internode (Martin *et al.* 2016). Therefore, *SD1* could also be considered as a candidate gene.

We next examined if any of these candidate genes had changes in protein sequence or gene expression patterns in the parental lines, *S. viridis* and *S. italica.* SD1 and the proteins encoded by four candidate genes have differences in the protein sequence between *S. viridis* and *S. italica* (Supplemental Table S4). One candidate Sevir.5G413600 (its *S. italica* ortholog, Seita.5G407900) is particularly interesting because the encoded protein contains four amino acid replacements between *S. viridis* and *S. italica*, which change the physicochemical property in a conserved C-terminal domain (Supplemental Figures S7 and S8). The protein is most similar to RIO2 kinase/ATPases. RIO2 proteins are widely conserved from archaea to eukaryotes and are involved in the maturation of small ribosome subunits during ribosome biogenesis through ATPase activity (LaRonde-LeBlanc and Wlodawer 2005; Knuppel *et al.* 2018). The protein has three domains (Ferreira-Cerca *et al.* 2012). While deletions in each of these domains render the protein non-functional and are lethal, a shorter truncation in the C-terminal domain is not lethal but leads to synthetic lethality with a non-essential ribosome factor called LTV1 (Ferreira-Cerca *et al.* 2012).

The SD1 protein sequence in *S. viridis* has two amino acid replacements (Supplemental Figure S9). The first substitution is a glutamate to aspartate change at position 157 of the *S. viridis* SD1 protein. This amino acid replacement occurs within a relatively conserved region across grass species (Supplemental Figure S9) but is a conservative replacement in the same physicochemical group and this change occurs in other grasses. The second substitution is an alanine to aspartate change at position 366 of the *S. viridis* SD1 protein. This amino acid replacement is a non-conservative replacement but the sequence nearby it is not conserved across grass species. Neither amino acid replacements are within the catalytic Fe^2^OG dioxygenase domain of SD1.

We also examined gene expression differences between *S. viridis* and *S. italica* for the thirteen candidate genes and the *SD1* gene based on the RNAseq data available at Phytozome12 (Supplemental Table S4). One candidate gene, Sevir.5G394900, has lower expression in most *S. viridis* tissues than its *S. italica* ortholog Seita.5G389700 (Supplemental Figure S10). This gene is annotated as a gene encoding a ribosomal protein belonging to the L1P family (Byrne 2009). The expression difference may be caused by polymorphisms in the promoter region of this gene. We therefore compared the 1kb upstream sequence flanking this gene between *S. viridis* and *S. italica*. We identified five SNPs and one insertion in *S. viridis,* including a SNP located at a predicted MYB transcription factor binding site (Supplemental Figure S11 and Supplemental Table S5).

## Discussion

The QTG-Finder we previously developed relies on known causal genes as a training set and cannot be extended to other plant species with few or no known causal genes. Since nearly all plant species, including important crops, do not currently have a sufficient set of cloned causal genes, this algorithm could not be applied to species beyond Arabidopsis and rice. Here, we have developed QTG-Finder2, which solves this problem by using the orthologs of causal genes to train models in other species.

Some orthologous genes have been repeatedly found to cause variation in similar traits across species. There are more than 100 examples showing that mutations occur at orthologous loci and cause similar phenotypic variation (Martin and Orgogozo 2013). Therefore, we posited that the orthologs of causal genes might determine similar trait variation as the causal genes. Why these genes become genetic hotspots of trait variation is still unknown but there are two theories (Martin and Orgogozo 2013). The first theory is mutational bias. The hotspot genes may be more prone to ectopic changes due to being in unstable chromosomal regions or structures like repeat-rich regions (Chan *et al.* 2010; Martin and Orgogozo 2013). The second theory is optimal pleiotropy (Kopp 2009). The hotspot genes may be able to generate variations in a trait without interfering with other traits. These hypotheses remain to be rigorously tested. In the meantime, given that many known causal genes are genetic hotspots of trait variation, we hypothesized, tested and showed that we can use an orthology approach to transfer the information about causal genes between species.

The major advantage of QTG-Finder2 over QTG-Finder is that it facilitates building models for species that have a limited number of known causal genes, which currently represents almost all plant species, including all major crops except rice. We have shown that Arabidopsis and rice models that were trained on orthologs of causal genes from other species have similar performance as models trained with known causal genes in Arabidopsis and rice. This result indicates that the orthologs derived from known causal genes in other species contain information that can be used to train models. As proof of concept, we applied QTG-Finder2 to train new models for *Setaria viridis* and *Sorghum bicolor*. The sorghum model has a 70% chance to recall a real causal gene (unseen during training) when the top 20% genes in a QTL are prioritized, which is a comparable performance to Arabidopsis and rice.

Sorghum is an important C4 crop with good drought resistance. There are 2605 sorghum QTLs identified by linkage mapping according to Sorghum QTL Atlas (Mace *et al.* 2019). For most of these QTLs, the causal genes have not been identified. The Sorghum model can be used to prioritize candidate genes and accelerate the discovery of causal genes in these QTLs. *Setaria viridis* has been developed as a model grass due to advantages like short life span, small plant stature and small diploid genome (Huang *et al.* 2016b). High-throughput phenotyping techniques have been developed for both underground and above-ground traits (Fahlgren *et al.* 2015; Rellan-Alvarez *et al.* 2015; Sebastian *et al.* 2016), which facilitate not only more QTL mapping studies but also faster phenotype screening for characterizing mutants of candidate genes. The Setaria model will play an important role in this pipeline by refining the candidates identified by QTL mapping for the downstream validation and functional analyses.

We have applied the Setaria model to prioritize genes in a height QTL and combined the results with published transcriptome, gene function and sequence data to generate a hypothesis for candidate genes. We prioritized 13 candidate genes including two strong candidates, a RIO2 kinase/ATPase gene (Sevir.5G413600) and an L1P ribosome protein gene (Sevir.5G394900). RIO2 has two domains, winged helix (wHTH) and kinase, which are conserved from archaea to eukaryotes, and a C-terminal extension domain that is conserved only in eukaryotes (LaRonde-LeBlanc and Wlodawer 2005; Ferreira-Cerca *et al.* 2012). While deletions in each of these domains render the protein non-functional and are lethal, a shorter truncation in the C-terminal domain was not lethal but led to synthetic lethality with a non-essential ribosome factor called LTV1 (Ferreira-Cerca *et al.* 2012). It is this region where there are four non-synonymous substitutions between *S. viridis* and *S. italica* (Supplementary Figure S7). RIO2 is found as a single-copy gene in most plants (Gao *et al.* 2018) but has not been functionally characterized in any plants to date.

The other candidate gene (Sevir.5G394900) encodes an L1P family ribosomal protein. L1P family ribosomal proteins are involved in binding and releasing de-acylated tRNA from the E site of ribosomes (Nikulin *et al.* 2003; Byrne 2009). Arabidopsis mutants of an L1P family ribosomal protein, *PGY1*, are not lethal and have subtle leaf phenotypes (Pinon *et al.* 2008). *PGY1* may function with proteins like *ASYMMETRIC LEAVES1 (AS1)* and *REVOLUTA (REV)* to affect plant development in different organs. For example, the *as1 pgy1* double mutant has ectopic leaf lamina outgrowth and the *rev pyg1* double mutant has inflorescence defects (Pinon *et al.* 2008). This candidate gene has higher expression in *S. italica* than *S. viridis* across leaf, shoot and root tissues (Supplemental Figure S10) and therefore may have a broad effect on development in Setaria. The expression difference of this gene is likely caused by a SNP in a putative MYB transcription factor binding site located in the promoter of this gene (Supplemental Table S5).

Though not prioritized as a top 20% gene, the *SD1* gene (Sevir.5G410400) is also a potential causal gene based on its function in other species and up-regulation in internode meristem zone relative to maturation zone. *SD1* gene encodes gibberellin20 oxidase2 in rice and a loss of function allele gives dwarf phenotype in rice (Spielmeyer *et al.* 2002). However, the rice *SD1* has three orthologs in *S. viridis*: Sevir.3G242400, Sevir.5G410400, Sevir.7G114500, and we do not know if they are functionally redundant.

In summary, we developed QTG-Finder2 algorithm by incorporating an orthology approach, which can be used to train models in species that have few or even no known causal genes. We have built new models using QTG-Finder2 for *S. bicolor* and *S. viridis* to accelerate the causal gene discovery in these cereal crops and models. The algorithm can also be potentially applied to other important crop species such as maize, barley and wheat to accelerate gene discovery and trait improvement.

## Acknowledgments

We thank Drs. L. Wang and S. Yang for sharing the rice gene presence/absence matrix. We acknowledge H. Nam for technical assistance. We thank all members in Rhee lab for useful discussions, especially Dr. J. Yen.

