## Supplementary material for "QTG-Finder2: a generalized machine-learning algorithm for prioritizing QTL causal genes in plants": Figure S10

### Slide 1
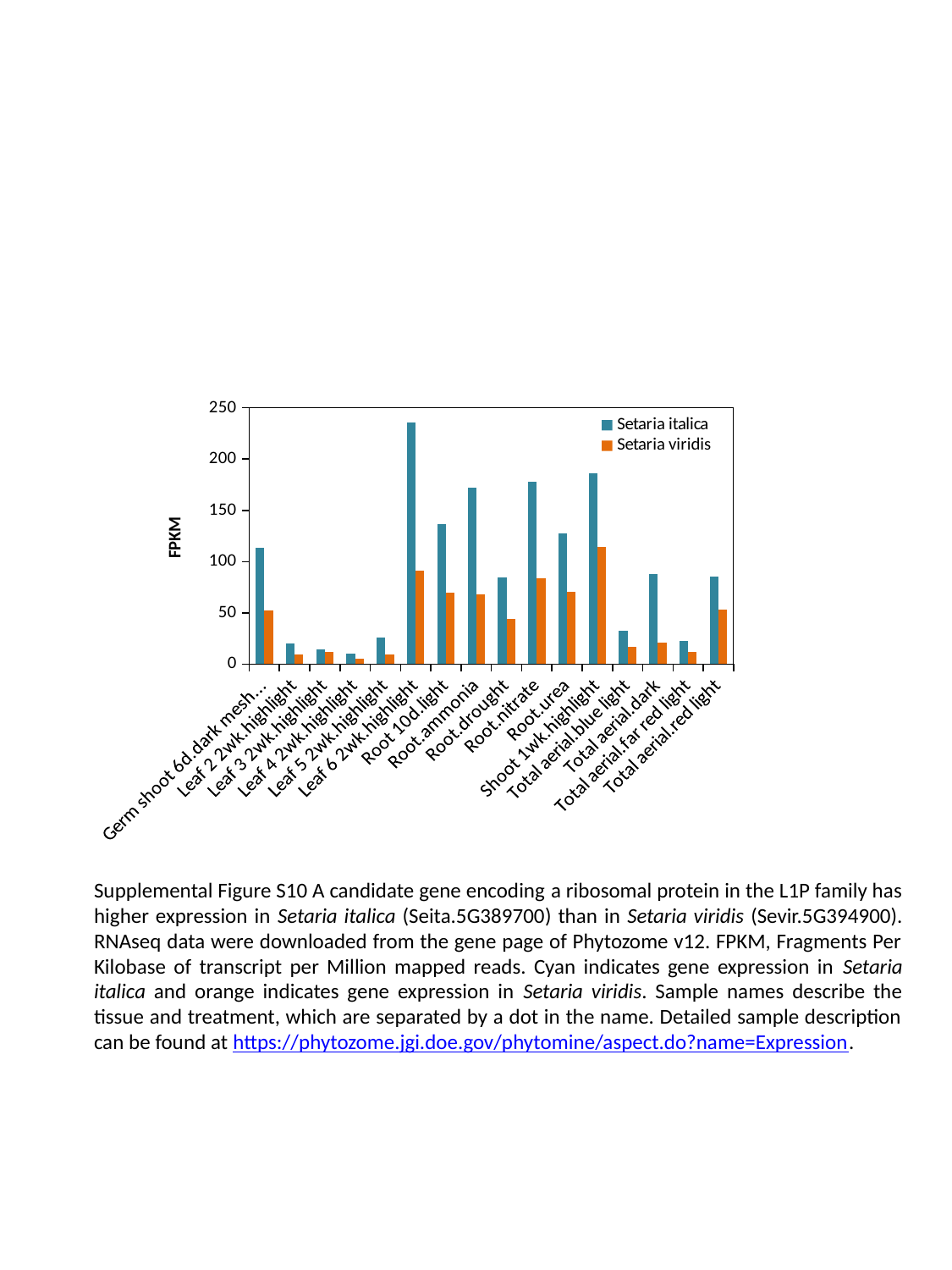

#### Chart
| Category | Setaria italica | Setaria viridis |
|---|---|---|
| Germ shoot 6d.dark mesh water | 113.736 | 52.466 |
| Leaf 2 2wk.highlight | 19.968 | 9.43 |
| Leaf 3 2wk.highlight | 14.618 | 12.082 |
| Leaf 4 2wk.highlight | 10.008 | 5.433 |
| Leaf 5 2wk.highlight | 25.64 | 9.734 |
| Leaf 6 2wk.highlight | 235.571 | 91.18299999999998 |
| Root 10d.light | 136.382 | 69.264 |
| Root.ammonia | 172.055 | 68.406 |
| Root.drought | 84.77299999999998 | 44.303 |
| Root.nitrate | 178.101 | 83.532 |
| Root.urea | 127.375 | 70.772 |
| Shoot 1wk.highlight | 186.297 | 114.415 |
| Total aerial.blue light | 32.416 | 16.834 |
| Total aerial.dark | 87.432 | 20.534 |
| Total aerial.far red light | 22.66 | 11.863 |
| Total aerial.red light | 85.62499999999999 | 53.183 |Supplemental Figure S10 A candidate gene encoding a ribosomal protein in the L1P family has higher expression in Setaria italica (Seita.5G389700) than in Setaria viridis (Sevir.5G394900). RNAseq data were downloaded from the gene page of Phytozome v12. FPKM, Fragments Per Kilobase of transcript per Million mapped reads. Cyan indicates gene expression in Setaria italica and orange indicates gene expression in Setaria viridis. Sample names describe the tissue and treatment, which are separated by a dot in the name. Detailed sample description can be found at https://phytozome.jgi.doe.gov/phytomine/aspect.do?name=Expression.
