## Supplementary material for "QTG-Finder2: a generalized machine-learning algorithm for prioritizing QTL causal genes in plants": Figure S1

### Slide 1
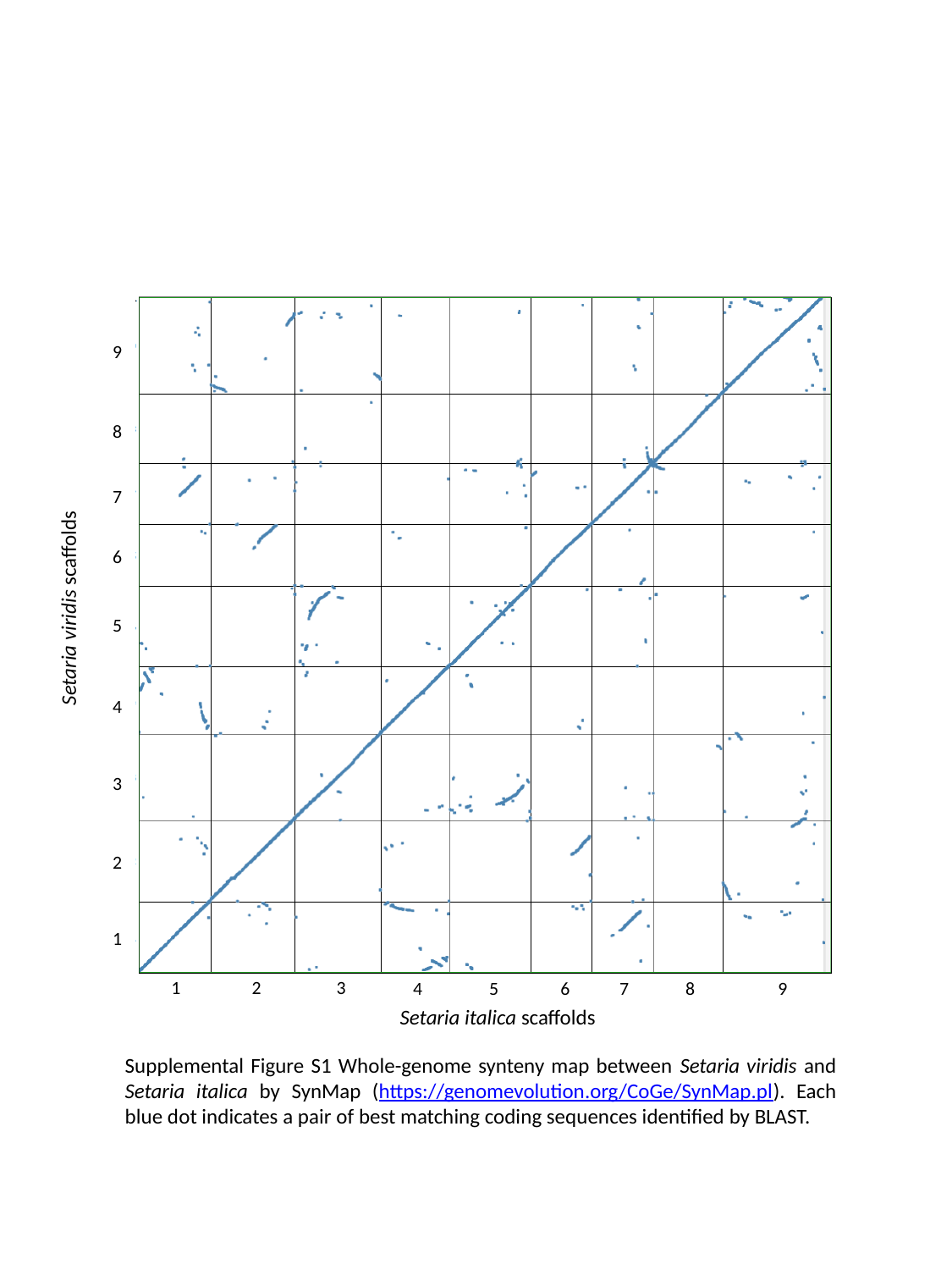

9
8
7
6
Setaria viridis scaffolds
5
4
3
2
1
1
2
3
4
5
6
7
8
9
Setaria italica scaffolds
Supplemental Figure S1 Whole-genome synteny map between Setaria viridis and Setaria italica by SynMap (https://genomevolution.org/CoGe/SynMap.pl). Each blue dot indicates a pair of best matching coding sequences identified by BLAST.
