## Supplementary material for "QTG-Finder2: a generalized machine-learning algorithm for prioritizing QTL causal genes in plants": Figure S2

### Slide 1
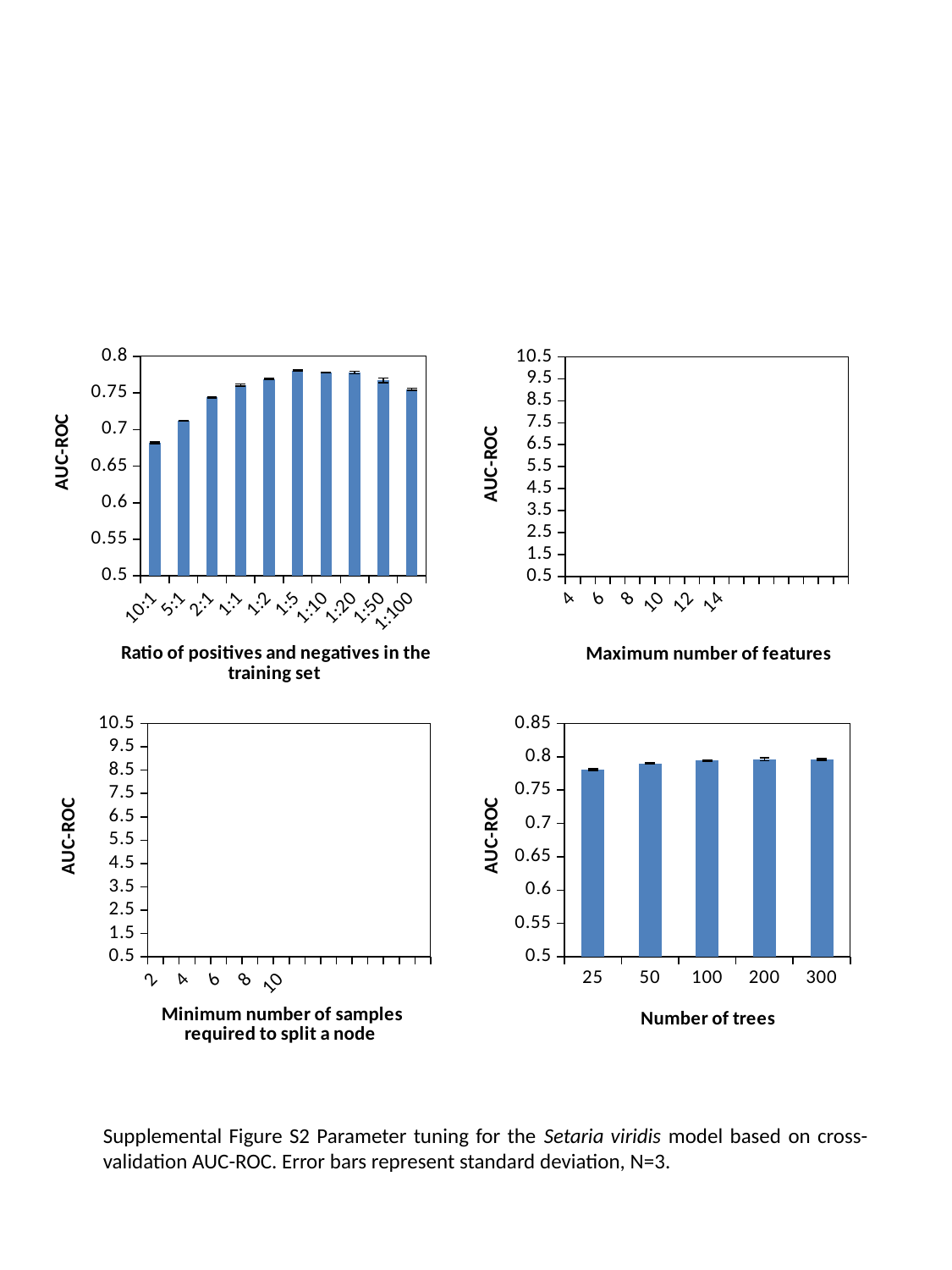

#### Chart
| Category | |
|---|---|
| 4.0 | 0.792148013580246 |
| 5.0 | 0.790778024902175 |
| 6.0 | 0.793603864917112 |
| 7.0 | 0.793699988900557 |
| 8.0 | 0.792295361373272 |
| 9.0 | 0.791961366292873 |
| 10.0 | 0.793444654014096 |
| 11.0 | 0.794367062095964 |
| 12.0 | 0.792084781259772 |
| 13.0 | 0.792350003525558 |
| 14.0 | 0.793783710036017 |
| 15.0 | 0.794846324088854 |
#### Chart
| Category | |
|---|---|
| 10:1 | 0.681569176753249 |
| 5:1 | 0.711526894164002 |
| 2:1 | 0.744078229113865 |
| 1:1 | 0.760609819328728 |
| 1:2 | 0.769102211488016 |
| 1:5 | 0.780178057403517 |
| 1:10 | 0.77786404129579 |
| 1:20 | 0.777882236511937 |
| 1:50 | 0.767312699113942 |
| 1:100 | 0.755052440873377 |
#### Chart
| Category | |
|---|---|
| 2.0 | 0.779078503174714 |
| 3.0 | 0.779162748938131 |
| 4.0 | 0.780234353791609 |
| 5.0 | 0.778575413504399 |
| 6.0 | 0.781110552316273 |
| 7.0 | 0.779206148084747 |
| 8.0 | 0.781101617619236 |
| 9.0 | 0.780099872282122 |
| 10.0 | 0.779861597723789 |
#### Chart
| Category | |
|---|---|
| 25.0 | 0.780887614782842 |
| 50.0 | 0.789915065968867 |
| 100.0 | 0.794351244852388 |
| 200.0 | 0.796131616564213 |
| 300.0 | 0.795896498055964 |Supplemental Figure S2 Parameter tuning for the Setaria viridis model based on cross-validation AUC-ROC. Error bars represent standard deviation, N=3.
