## Supplementary material for "QTG-Finder2: a generalized machine-learning algorithm for prioritizing QTL causal genes in plants": Figure S3

### Slide 1
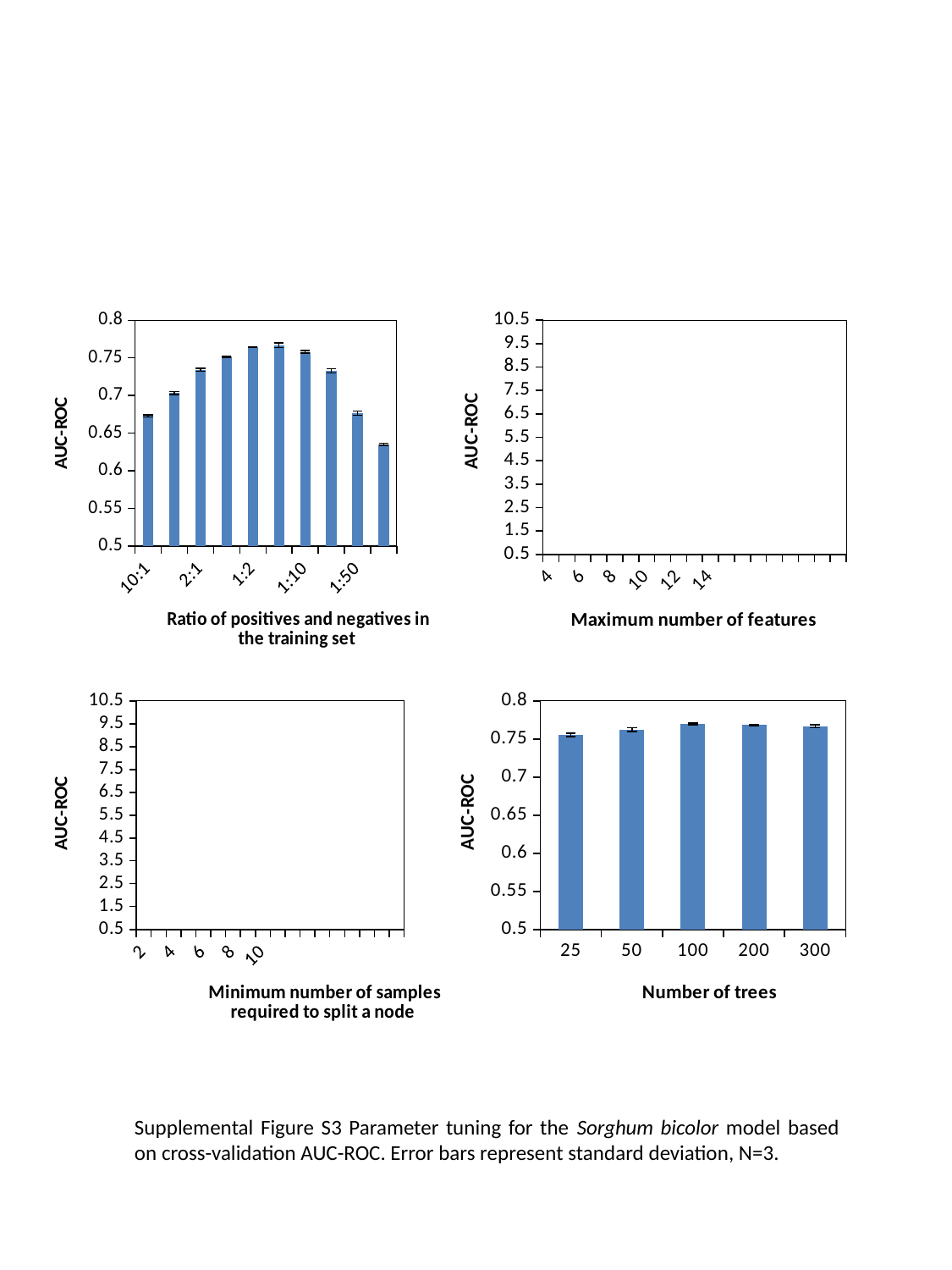

#### Chart
| Category | |
|---|---|
| 4.0 | 0.76708958562403 |
| 5.0 | 0.767732555614965 |
| 6.0 | 0.769303269784814 |
| 7.0 | 0.769723673106924 |
| 8.0 | 0.766754796111058 |
| 9.0 | 0.767008443543959 |
| 10.0 | 0.770125494246024 |
| 11.0 | 0.770666704301722 |
| 12.0 | 0.769108295737903 |
| 13.0 | 0.765891696121141 |
| 14.0 | 0.769857593316626 |
| 15.0 | 0.771984291355816 |
#### Chart
| Category | |
|---|---|
| 10:1 | 0.672890595254464 |
| 5:1 | 0.703357984816144 |
| 2:1 | 0.734065986465718 |
| 1:1 | 0.751697985631901 |
| 1:2 | 0.764527859038557 |
| 1:5 | 0.766903338046511 |
| 1:10 | 0.75787278007328 |
| 1:20 | 0.732741530530862 |
| 1:50 | 0.676510542896524 |
| 1:100 | 0.635193718538339 |
#### Chart
| Category | |
|---|---|
| 2.0 | 0.764266844772825 |
| 3.0 | 0.769078128479887 |
| 4.0 | 0.772123252406033 |
| 5.0 | 0.772404683777511 |
| 6.0 | 0.7763839717407 |
| 7.0 | 0.775322792302952 |
| 8.0 | 0.777768660090333 |
| 9.0 | 0.776572580048791 |
| 10.0 | 0.777387457876289 |
#### Chart
| Category | |
|---|---|
| 25.0 | 0.755202715105518 |
| 50.0 | 0.762320508791995 |
| 100.0 | 0.76976638085051 |
| 200.0 | 0.768119053175545 |
| 300.0 | 0.766526994013835 |Supplemental Figure S3 Parameter tuning for the Sorghum bicolor model based on cross-validation AUC-ROC. Error bars represent standard deviation, N=3.
