## Supplementary material for "QTG-Finder2: a generalized machine-learning algorithm for prioritizing QTL causal genes in plants": Figure S4

### Slide 1
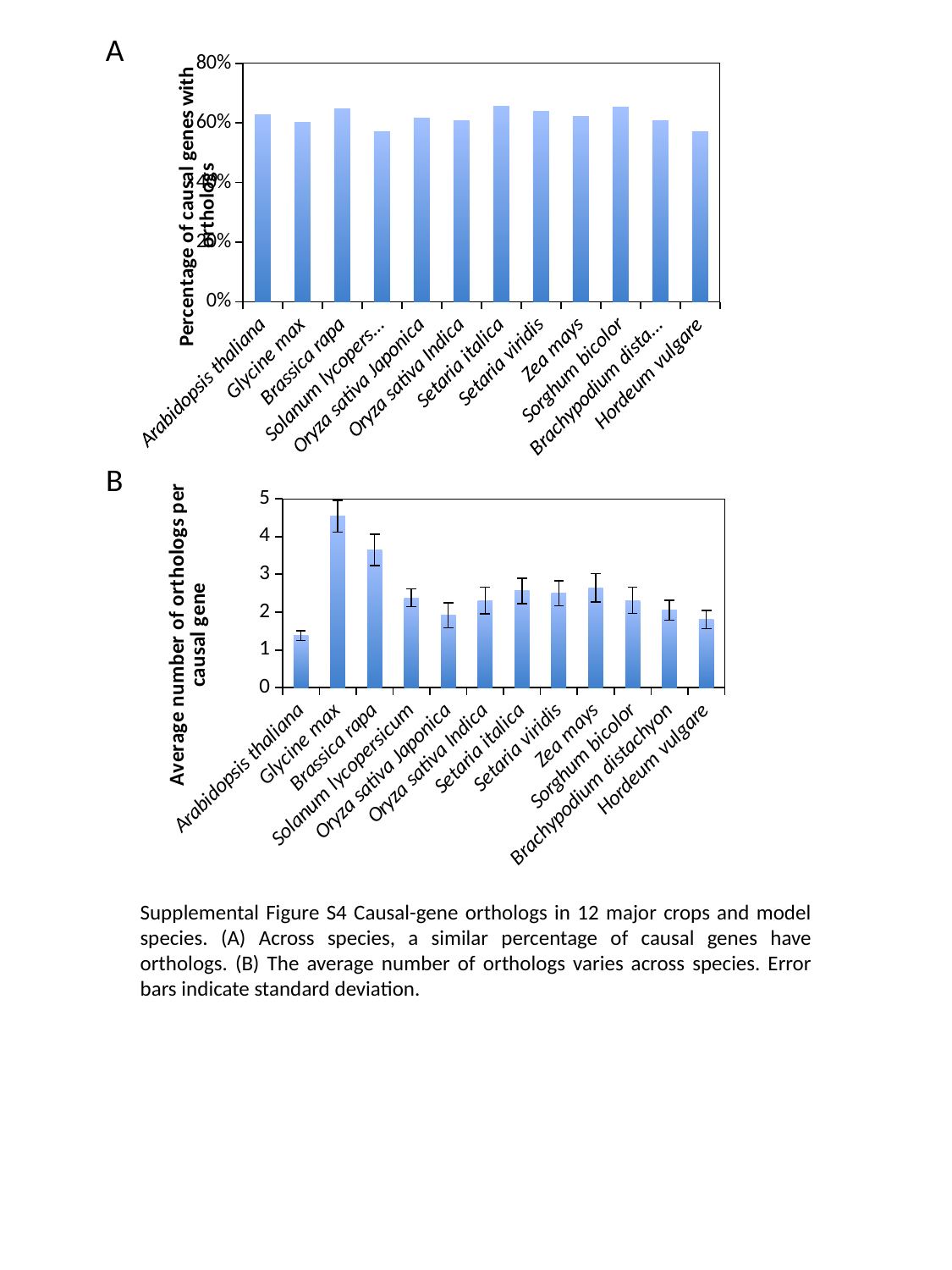

A
#### Chart
| Category | No ortholog percentage |
|---|---|
| Arabidopsis thaliana | 0.629032258064517 |
| Glycine max | 0.603238866396762 |
| Brassica rapa | 0.648 |
| Solanum lycopersicum | 0.569620253164557 |
| Oryza sativa Japonica | 0.61576354679803 |
| Oryza sativa Indica | 0.608695652173913 |
| Setaria italica | 0.654761904761905 |
| Setaria viridis | 0.63774062816616 |
| Zea mays | 0.62280701754386 |
| Sorghum bicolor | 0.653225806451613 |
| Brachypodium distachyon | 0.608695652173913 |
| Hordeum vulgare | 0.571428571428572 |B
#### Chart
| Category | Avg_orth |
|---|---|
| Arabidopsis thaliana | 1.375 |
| Glycine max | 4.535483870967735 |
| Brassica rapa | 3.64242424242424 |
| Solanum lycopersicum | 2.377483443708598 |
| Oryza sativa Japonica | 1.91428571428571 |
| Oryza sativa Indica | 2.3051948051948 |
| Setaria italica | 2.56024096385542 |
| Setaria viridis | 2.496411176621378 |
| Zea mays | 2.64071856287425 |
| Sorghum bicolor | 2.31137724550898 |
| Brachypodium distachyon | 2.05194805194805 |
| Hordeum vulgare | 1.80132450331125 |Supplemental Figure S4 Causal-gene orthologs in 12 major crops and model species. (A) Across species, a similar percentage of causal genes have orthologs. (B) The average number of orthologs varies across species. Error bars indicate standard deviation.
