## Supplementary material for "QTG-Finder2: a generalized machine-learning algorithm for prioritizing QTL causal genes in plants": Figure S5

### Slide 1
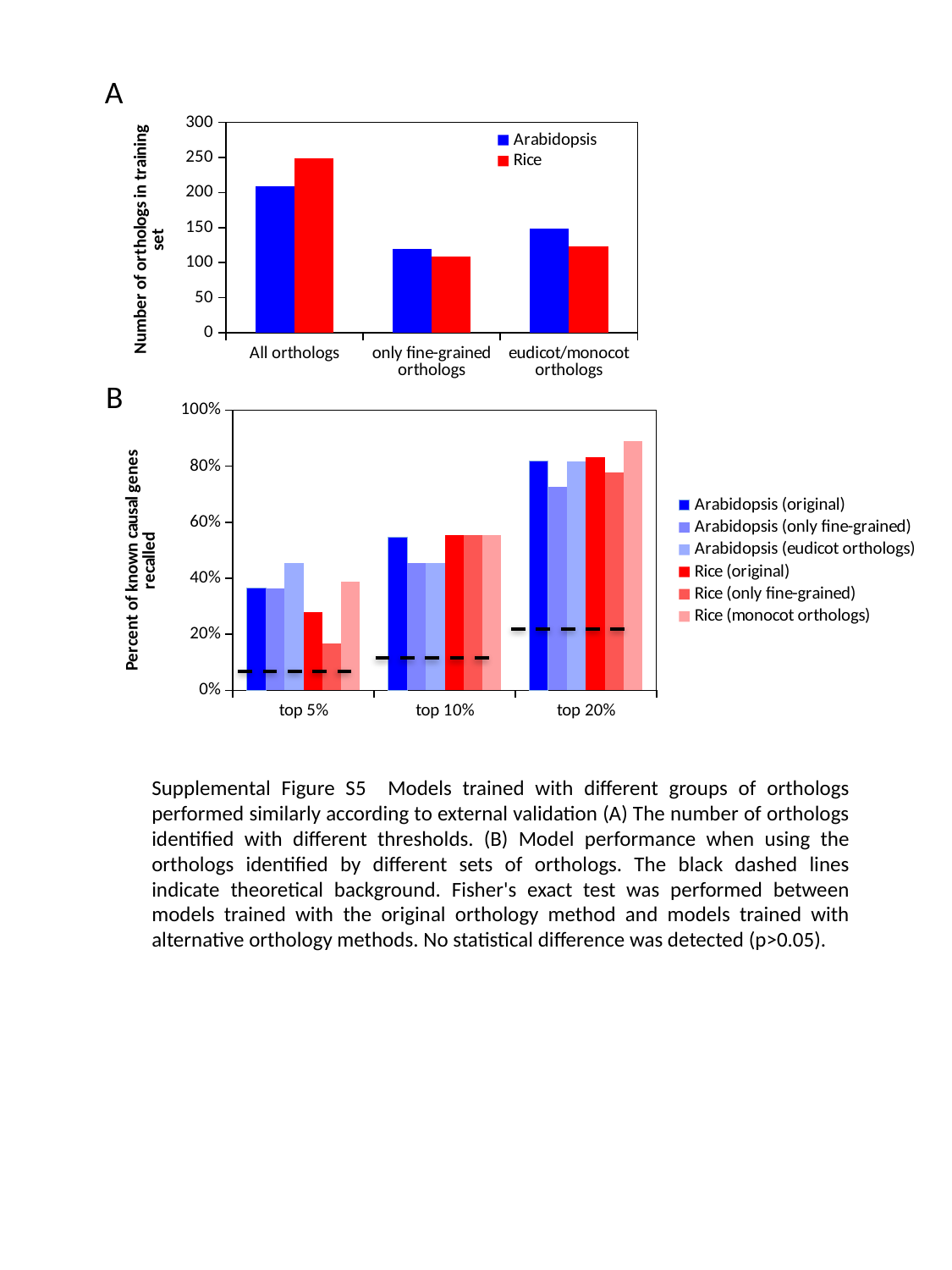

A
#### Chart
| Category | Arabidopsis | Rice |
|---|---|---|
| All orthologs | 209.0 | 249.0 |
| only fine-grained orthologs | 120.0 | 109.0 |
| eudicot/monocot orthologs | 148.0 | 123.0 |B
#### Chart
| Category | Arabidopsis (original) | Arabidopsis (only fine-grained) | Arabidopsis (eudicot orthologs) | Rice (original) | Rice (only fine-grained) | Rice (monocot orthologs) |
|---|---|---|---|---|---|---|
| top 5% | 0.363636363636364 | 0.363636363636364 | 0.454545454545454 | 0.277777777777778 | 0.166666666666667 | 0.388888888888889 |
| top 10% | 0.545454545454545 | 0.454545454545454 | 0.454545454545454 | 0.555555555555556 | 0.555555555555556 | 0.555555555555556 |
| top 20% | 0.818181818181818 | 0.727272727272727 | 0.818181818181818 | 0.833333333333333 | 0.777777777777778 | 0.888888888888889 |Supplemental Figure S5 Models trained with different groups of orthologs performed similarly according to external validation (A) The number of orthologs identified with different thresholds. (B) Model performance when using the orthologs identified by different sets of orthologs. The black dashed lines indicate theoretical background. Fisher's exact test was performed between models trained with the original orthology method and models trained with alternative orthology methods. No statistical difference was detected (p>0.05).
