## Supplementary material for "QTG-Finder2: a generalized machine-learning algorithm for prioritizing QTL causal genes in plants": Figure S11

### Slide 1
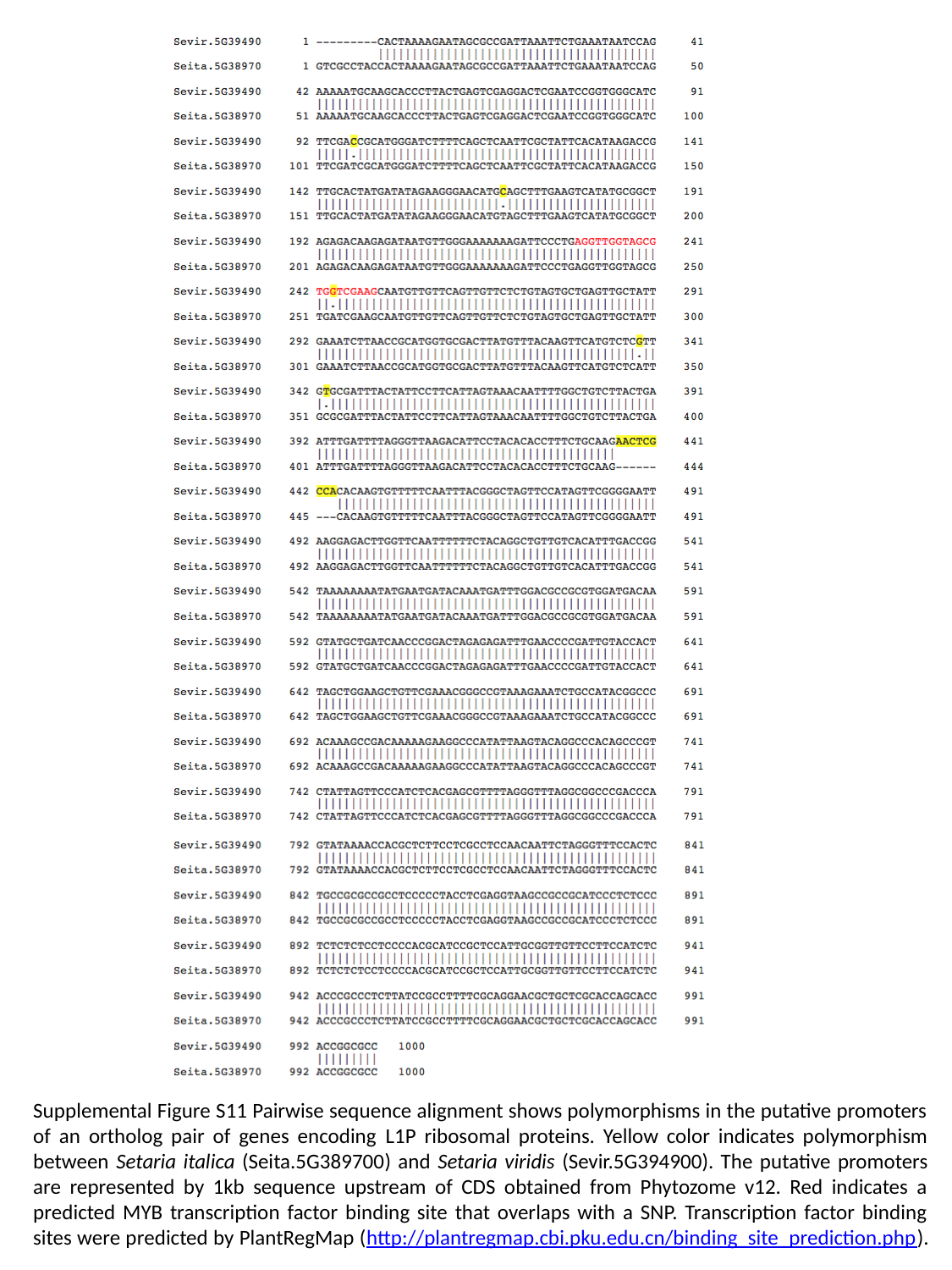

Supplemental Figure S11 Pairwise sequence alignment shows polymorphisms in the putative promoters of an ortholog pair of genes encoding L1P ribosomal proteins. Yellow color indicates polymorphism between Setaria italica (Seita.5G389700) and Setaria viridis (Sevir.5G394900). The putative promoters are represented by 1kb sequence upstream of CDS obtained from Phytozome v12. Red indicates a predicted MYB transcription factor binding site that overlaps with a SNP. Transcription factor binding sites were predicted by PlantRegMap (http://plantregmap.cbi.pku.edu.cn/binding_site_prediction.php).
