## Supplementary material for "QTG-Finder2: a generalized machine-learning algorithm for prioritizing QTL causal genes in plants": Figure S8

### Slide 1
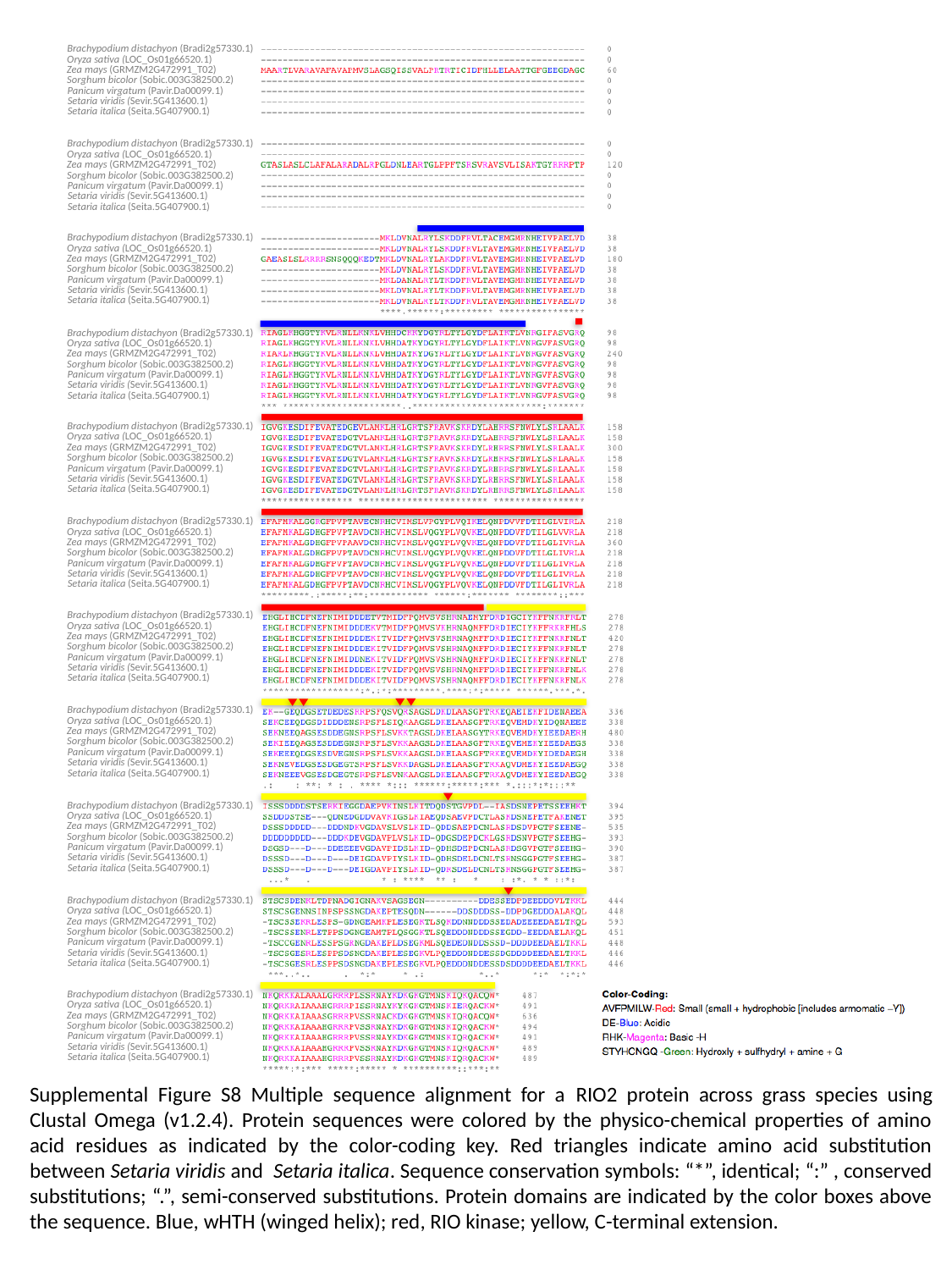

Brachypodium distachyon (Bradi2g57330.1)
Oryza sativa (LOC_Os01g66520.1)
Zea mays (GRMZM2G472991_T02)
Sorghum bicolor (Sobic.003G382500.2)
Panicum virgatum (Pavir.Da00099.1)
Setaria viridis (Sevir.5G413600.1)
Setaria italica (Seita.5G407900.1)
Brachypodium distachyon (Bradi2g57330.1)
Oryza sativa (LOC_Os01g66520.1)
Zea mays (GRMZM2G472991_T02)
Sorghum bicolor (Sobic.003G382500.2)
Panicum virgatum (Pavir.Da00099.1)
Setaria viridis (Sevir.5G413600.1)
Setaria italica (Seita.5G407900.1)
Brachypodium distachyon (Bradi2g57330.1)
Oryza sativa (LOC_Os01g66520.1)
Zea mays (GRMZM2G472991_T02)
Sorghum bicolor (Sobic.003G382500.2)
Panicum virgatum (Pavir.Da00099.1)
Setaria viridis (Sevir.5G413600.1)
Setaria italica (Seita.5G407900.1)
Brachypodium distachyon (Bradi2g57330.1)
Oryza sativa (LOC_Os01g66520.1)
Zea mays (GRMZM2G472991_T02)
Sorghum bicolor (Sobic.003G382500.2)
Panicum virgatum (Pavir.Da00099.1)
Setaria viridis (Sevir.5G413600.1)
Setaria italica (Seita.5G407900.1)
Brachypodium distachyon (Bradi2g57330.1)
Oryza sativa (LOC_Os01g66520.1)
Zea mays (GRMZM2G472991_T02)
Sorghum bicolor (Sobic.003G382500.2)
Panicum virgatum (Pavir.Da00099.1)
Setaria viridis (Sevir.5G413600.1)
Setaria italica (Seita.5G407900.1)
Brachypodium distachyon (Bradi2g57330.1)
Oryza sativa (LOC_Os01g66520.1)
Zea mays (GRMZM2G472991_T02)
Sorghum bicolor (Sobic.003G382500.2)
Panicum virgatum (Pavir.Da00099.1)
Setaria viridis (Sevir.5G413600.1)
Setaria italica (Seita.5G407900.1)
Brachypodium distachyon (Bradi2g57330.1)
Oryza sativa (LOC_Os01g66520.1)
Zea mays (GRMZM2G472991_T02)
Sorghum bicolor (Sobic.003G382500.2)
Panicum virgatum (Pavir.Da00099.1)
Setaria viridis (Sevir.5G413600.1)
Setaria italica (Seita.5G407900.1)
Brachypodium distachyon (Bradi2g57330.1)
Oryza sativa (LOC_Os01g66520.1)
Zea mays (GRMZM2G472991_T02)
Sorghum bicolor (Sobic.003G382500.2)
Panicum virgatum (Pavir.Da00099.1)
Setaria viridis (Sevir.5G413600.1)
Setaria italica (Seita.5G407900.1)
Brachypodium distachyon (Bradi2g57330.1)
Oryza sativa (LOC_Os01g66520.1)
Zea mays (GRMZM2G472991_T02)
Sorghum bicolor (Sobic.003G382500.2)
Panicum virgatum (Pavir.Da00099.1)
Setaria viridis (Sevir.5G413600.1)
Setaria italica (Seita.5G407900.1)
Brachypodium distachyon (Bradi2g57330.1)
Oryza sativa (LOC_Os01g66520.1)
Zea mays (GRMZM2G472991_T02)
Sorghum bicolor (Sobic.003G382500.2)
Panicum virgatum (Pavir.Da00099.1)
Setaria viridis (Sevir.5G413600.1)
Setaria italica (Seita.5G407900.1)
Brachypodium distachyon (Bradi2g57330.1)
Oryza sativa (LOC_Os01g66520.1)
Zea mays (GRMZM2G472991_T02)
Sorghum bicolor (Sobic.003G382500.2)
Panicum virgatum (Pavir.Da00099.1)
Setaria viridis (Sevir.5G413600.1)
Setaria italica (Seita.5G407900.1)
Supplemental Figure S8 Multiple sequence alignment for a RIO2 protein across grass species using Clustal Omega (v1.2.4). Protein sequences were colored by the physico-chemical properties of amino acid residues as indicated by the color-coding key. Red triangles indicate amino acid substitution between Setaria viridis and Setaria italica. Sequence conservation symbols: “*”, identical; “:” , conserved substitutions; “.”, semi-conserved substitutions. Protein domains are indicated by the color boxes above the sequence. Blue, wHTH (winged helix); red, RIO kinase; yellow, C-terminal extension.
