## Supplementary material for "QTG-Finder2: a generalized machine-learning algorithm for prioritizing QTL causal genes in plants": Figure S9

### Slide 1
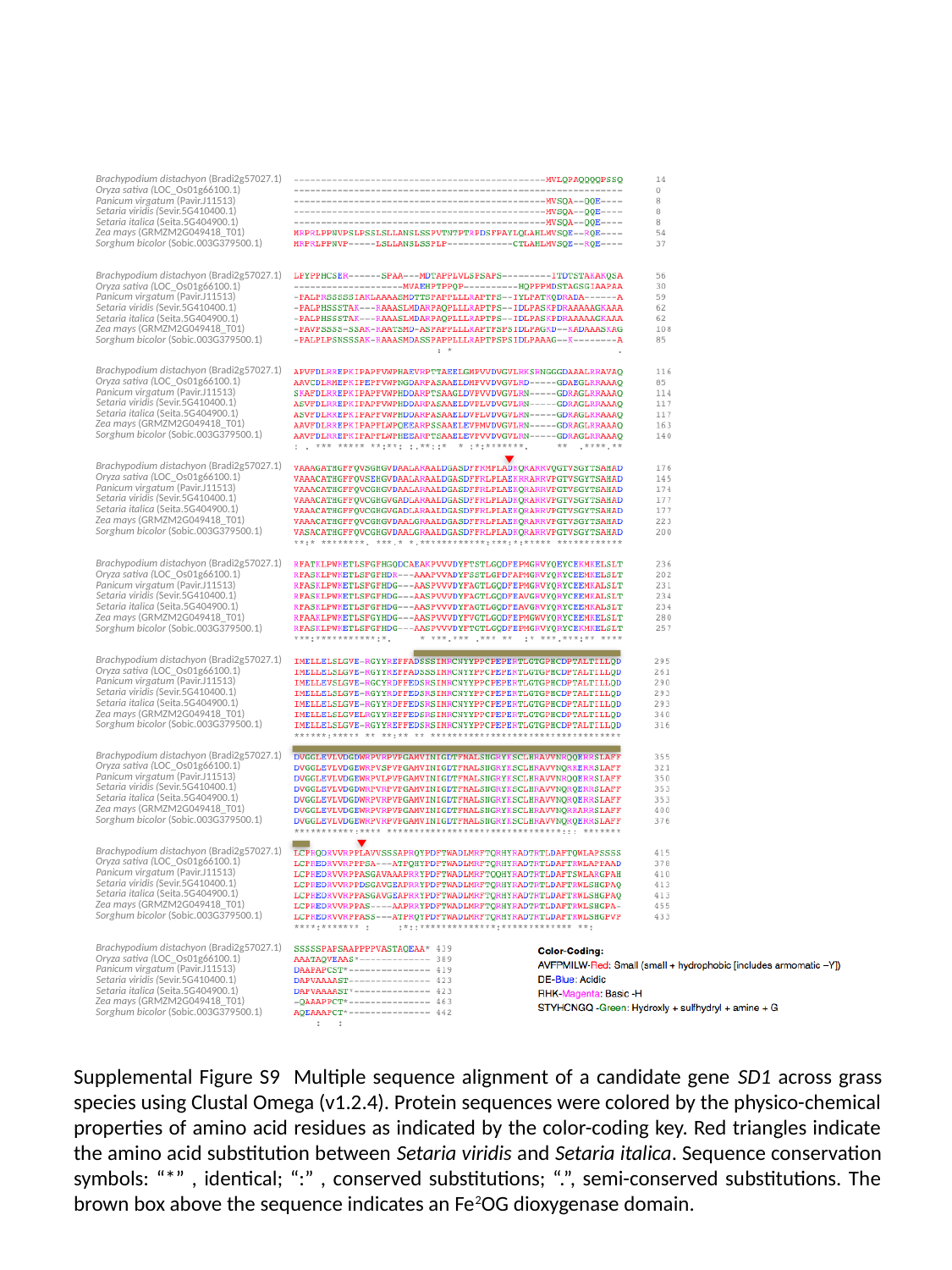

Brachypodium distachyon (Bradi2g57027.1)
Oryza sativa (LOC_Os01g66100.1)
Panicum virgatum (Pavir.J11513)
Setaria viridis (Sevir.5G410400.1)
Setaria italica (Seita.5G404900.1)
Zea mays (GRMZM2G049418_T01)
Sorghum bicolor (Sobic.003G379500.1)
Brachypodium distachyon (Bradi2g57027.1)
Oryza sativa (LOC_Os01g66100.1)
Panicum virgatum (Pavir.J11513)
Setaria viridis (Sevir.5G410400.1)
Setaria italica (Seita.5G404900.1)
Zea mays (GRMZM2G049418_T01)
Sorghum bicolor (Sobic.003G379500.1)
Brachypodium distachyon (Bradi2g57027.1)
Oryza sativa (LOC_Os01g66100.1)
Panicum virgatum (Pavir.J11513)
Setaria viridis (Sevir.5G410400.1)
Setaria italica (Seita.5G404900.1)
Zea mays (GRMZM2G049418_T01)
Sorghum bicolor (Sobic.003G379500.1)
Brachypodium distachyon (Bradi2g57027.1)
Oryza sativa (LOC_Os01g66100.1)
Panicum virgatum (Pavir.J11513)
Setaria viridis (Sevir.5G410400.1)
Setaria italica (Seita.5G404900.1)
Zea mays (GRMZM2G049418_T01)
Sorghum bicolor (Sobic.003G379500.1)
Brachypodium distachyon (Bradi2g57027.1)
Oryza sativa (LOC_Os01g66100.1)
Panicum virgatum (Pavir.J11513)
Setaria viridis (Sevir.5G410400.1)
Setaria italica (Seita.5G404900.1)
Zea mays (GRMZM2G049418_T01)
Sorghum bicolor (Sobic.003G379500.1)
Brachypodium distachyon (Bradi2g57027.1)
Oryza sativa (LOC_Os01g66100.1)
Panicum virgatum (Pavir.J11513)
Setaria viridis (Sevir.5G410400.1)
Setaria italica (Seita.5G404900.1)
Zea mays (GRMZM2G049418_T01)
Sorghum bicolor (Sobic.003G379500.1)
Brachypodium distachyon (Bradi2g57027.1)
Oryza sativa (LOC_Os01g66100.1)
Panicum virgatum (Pavir.J11513)
Setaria viridis (Sevir.5G410400.1)
Setaria italica (Seita.5G404900.1)
Zea mays (GRMZM2G049418_T01)
Sorghum bicolor (Sobic.003G379500.1)
Brachypodium distachyon (Bradi2g57027.1)
Oryza sativa (LOC_Os01g66100.1)
Panicum virgatum (Pavir.J11513)
Setaria viridis (Sevir.5G410400.1)
Setaria italica (Seita.5G404900.1)
Zea mays (GRMZM2G049418_T01)
Sorghum bicolor (Sobic.003G379500.1)
Brachypodium distachyon (Bradi2g57027.1)
Oryza sativa (LOC_Os01g66100.1)
Panicum virgatum (Pavir.J11513)
Setaria viridis (Sevir.5G410400.1)
Setaria italica (Seita.5G404900.1)
Zea mays (GRMZM2G049418_T01)
Sorghum bicolor (Sobic.003G379500.1)
Supplemental Figure S9 Multiple sequence alignment of a candidate gene SD1 across grass species using Clustal Omega (v1.2.4). Protein sequences were colored by the physico-chemical properties of amino acid residues as indicated by the color-coding key. Red triangles indicate the amino acid substitution between Setaria viridis and Setaria italica. Sequence conservation symbols: “*” , identical; “:” , conserved substitutions; “.”, semi-conserved substitutions. The brown box above the sequence indicates an Fe^2^OG dioxygenase domain.
