## Supplementary material for "QTG-Finder2: a generalized machine-learning algorithm for prioritizing QTL causal genes in plants": Figure S6

### Slide 1
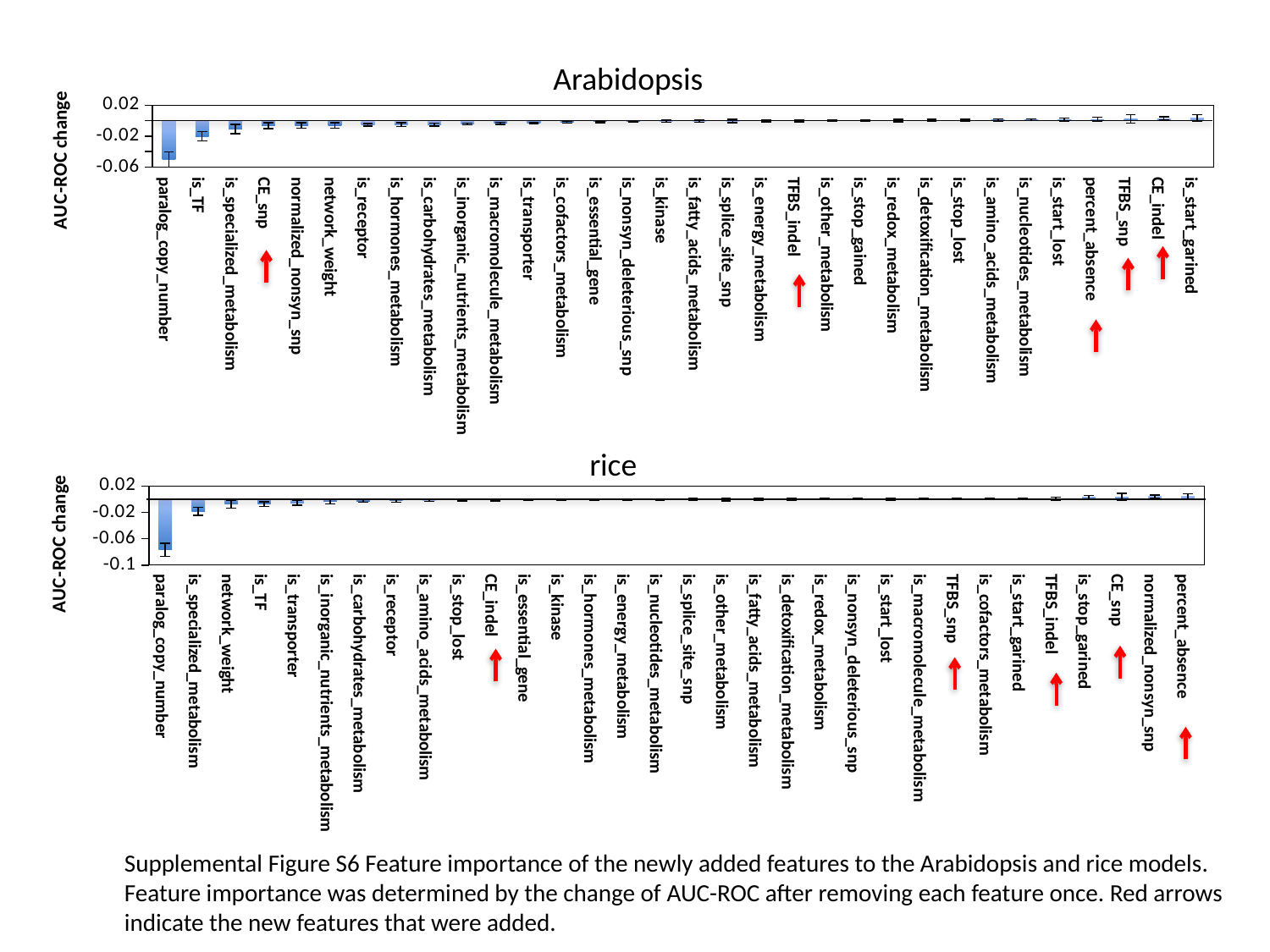

Arabidopsis
#### Chart
| Category | ROC-AUC change |
|---|---|
| paralog_copy_number | -0.0502608287319378 |
| is_TF | -0.0200829245845891 |
| is_specialized_metabolism | -0.010783713453004 |
| CE_snp | -0.00639037511134403 |
| normalized_nonsyn_SNP | -0.00609041630638639 |
| network_weight | -0.00608749554605149 |
| is_receptor | -0.00528463603954592 |
| is_hormones_metabolism | -0.00519365802230073 |
| is_carbohydrates_metabolism | -0.00511619591730318 |
| is_inorganic_nutrients_metabolism | -0.00436643748981735 |
| is_macromolecule_metabolism | -0.00327511786560747 |
| is_transporter | -0.00280722278090957 |
| is_cofactors_metabolism | -0.00205774308913862 |
| is_essential_gene | -0.00124239325399448 |
| is_nonsyn_deleterious | -0.00108034137033754 |
| is_kinase | -0.000868147538313362 |
| is_fatty_acids_lipids_metabolism | -0.000631912895658185 |
| is_splice_site_SNP | -0.000296366603970779 |
| is_energy_metabolism | -0.000192824246813757 |
| TFBS_indel | 2.77924917515845e-05 |
| is_other_metabolism | 6.82490236273422e-05 |
| is_stop_gained | 0.000210574482354248 |
| is_redox_metabolism | 0.000256522815914653 |
| is_detoxification_metabolism | 0.000617628402915577 |
| is_stop_lost | 0.000629632368546601 |
| is_amino_acids_metabolism | 0.00110648377637128 |
| is_nucleotides_metabolism | 0.001296583389277 |
| is_start_lost | 0.00132085596166992 |
| percent_absence | 0.00180998913557314 |
| TFBS_snp | 0.00249916343351185 |
| CE_indel | 0.00290814084624309 |
| is_start_gained | 0.00363177623702222 || paralog\_copy\_number | is\_TF | is\_specialized\_metabolism | CE\_snp | normalized\_nonsyn\_snp | network\_weight | is\_receptor | is\_hormones\_metabolism | is\_carbohydrates\_metabolism | is\_inorganic\_nutrients\_metabolism | is\_macromolecule\_metabolism | is\_transporter | is\_cofactors\_metabolism | is\_essential\_gene | is\_nonsyn\_deleterious\_snp | is\_kinase | is\_fatty\_acids\_metabolism | is\_splice\_site\_snp | is\_energy\_metabolism | TFBS\_indel | is\_other\_metabolism | is\_stop\_gained | is\_redox\_metabolism | is\_detoxification\_metabolism | is\_stop\_lost | is\_amino\_acids\_metabolism | is\_nucleotides\_metabolism | is\_start\_lost | percent\_absence | TFBS\_snp | CE\_indel | is\_start\_garined |
| --- | --- | --- | --- | --- | --- | --- | --- | --- | --- | --- | --- | --- | --- | --- | --- | --- | --- | --- | --- | --- | --- | --- | --- | --- | --- | --- | --- | --- | --- | --- | --- |
rice
#### Chart
| Category | ROC-AUC change |
|---|---|
| paralog_copy_number | -0.076898743081864 |
| is_specialized_metabolism | -0.0181860193595261 |
| network_weight | -0.00749620604155675 |
| is_TF | -0.00734355979184599 |
| is_transporter | -0.00543878227433013 |
| is_inorganic_nutrients_metabolism | -0.00310478709445138 |
| is_carbohydrates_metabolism | -0.00245463617122495 |
| is_receptor | -0.00171944790087463 |
| is_amino_acids_metabolism | -0.0011903649190615 |
| is_stop_lost | -0.00107938195270489 |
| CE_indel | -0.000922002376787447 |
| is_essential_gene | -0.000859419417187281 |
| is_kinase | -0.000686685457170624 |
| is_hormones_metabolism | -0.000686295910130038 |
| is_energy_metabolism | -0.000373201154056181 |
| is_nucleotides_metabolism | -0.000354405189596927 |
| is_splice_site_SNP | -9.3284548752835e-05 |
| is_other_metabolism | -8.55893905317221e-05 |
| is_fatty_acids_lipids_metabolism | 0.000201741270951919 |
| is_detoxification_metabolism | 0.00031530182858994 |
| is_redox_metabolism | 0.000360117216437596 |
| is_nonsyn_deleterious | 0.000438724796612523 |
| is_start_lost | 0.000440156357628765 |
| is_macromolecule_metabolism | 0.000630223877088249 |
| TFBS_snp | 0.000718737748900921 |
| is_cofactors_metabolism | 0.000808503634504143 |
| is_start_gained | 0.000852377424499051 |
| TFBS_indel | 0.00112375423610532 |
| is_stop_gained | 0.00314523448165116 |
| CE_snp | 0.00349819199305844 |
| normalized_nonsyn_SNP | 0.00405192644509232 |
| percent_absence | 0.00408592816928132 || paralog\_copy\_number | is\_specialized\_metabolism | network\_weight | is\_TF | is\_transporter | is\_inorganic\_nutrients\_metabolism | is\_carbohydrates\_metabolism | is\_receptor | is\_amino\_acids\_metabolism | is\_stop\_lost | CE\_indel | is\_essential\_gene | is\_kinase | is\_hormones\_metabolism | is\_energy\_metabolism | is\_nucleotides\_metabolism | is\_splice\_site\_snp | is\_other\_metabolism | is\_fatty\_acids\_metabolism | is\_detoxification\_metabolism | is\_redox\_metabolism | is\_nonsyn\_deleterious\_snp | is\_start\_lost | is\_macromolecule\_metabolism | TFBS\_snp | is\_cofactors\_metabolism | is\_start\_garined | TFBS\_indel | is\_stop\_garined | CE\_snp | normalized\_nonsyn\_snp | percent\_absence |
| --- | --- | --- | --- | --- | --- | --- | --- | --- | --- | --- | --- | --- | --- | --- | --- | --- | --- | --- | --- | --- | --- | --- | --- | --- | --- | --- | --- | --- | --- | --- | --- |
Supplemental Figure S6 Feature importance of the newly added features to the Arabidopsis and rice models. Feature importance was determined by the change of AUC-ROC after removing each feature once. Red arrows indicate the new features that were added.
